# The CTPase activity of ParB acts as a timing mechanism to control the dynamics and function of prokaryotic DNA partition complexes

**DOI:** 10.1101/2021.05.05.442810

**Authors:** Manuel Osorio-Valeriano, Florian Altegoer, Chandan K. Das, Wieland Steinchen, Gaël Panis, Lara Connolley, Giacomo Giacomelli, Helge Feddersen, Laura Corrales-Guerrero, Pietro Giammarinaro, Juri Hanßmann, Marc Bramkamp, Patrick H. Viollier, Seán Murray, Lars V. Schäfer, Gert Bange, Martin Thanbichler

**Affiliations:** Department of Biology, University of Marburg, 35043 Marburg, Germany; Max Planck Institute for Terrestrial Microbiology, 35043 Marburg, Germany; Department of Chemistry, University of Marburg, 35043 Marburg, Germany; Center for Synthetic Microbiology, 35043 Marburg, Germany; Theoretical Chemistry, Ruhr University Bochum, 44801 Bochum, Germany; Department of Microbiology and Molecular Medicine, University of Geneva, 1211 Geneva, Switzerland; Department of Systems & Synthetic Microbiology, Max Planck Institute for Terrestrial Microbiology, 35043 Marburg, Germany; Institute for General Microbiology, Christian Albrechts University, 24118 Kiel, Germany

**Keywords:** Bacterial cell biology, DNA segregation, ParAB*S* system, ParA, *parS*, partition complex, molecular switch, nucleotide-binding protein, DNA-sliding clamp, ParB/sulfiredoxin domain, ParB-like nuclease domain, CTPase, NTPase, nucleotide hydrolysis

## Abstract

DNA partitioning CTPases of the ParB family mediate the segregation of bacterial chromosomes and low-copy number plasmids. They act as DNA-sliding clamps that are loaded at *parS* motifs in the centromeric region of target DNA molecules and then spread laterally to form large nucleoprotein complexes that serve as docking points for the DNA segregation machinery. Here, we identify conformational changes that underlie the CTP- and *parS*-dependent closure of ParB clamps. Moreover, we solve crystal structures of ParB in the pre- and post-hydrolysis state and provide insights into the catalytic mechanism underlying nucleotide hydrolysis. The characterization of CTPase-deficient ParB variants reveals that CTP hydrolysis serves as a timing mechanism to control the sliding time of ParB. Hyperstable clamps are trapped on the DNA, leading to excessing spreading and severe chromosome segregation defects *in vivo*. These findings clarify the role of the ParB CTPase cycle in partition complex dynamics and function and thus complete our understanding of this prototypic CTP-dependent molecular switch.

## INTRODUCTION

The regulation of cellular processes critically depends on proteins that use nucleotide binding and hydrolysis to switch between different activity states. Although a large number of such proteins have been reported to date, their repertoire of nucleotide cofactors has long been limited to the purine nucleotides guanosine and adenosine triphosphate (Bange and Sinning, 2013; Leipe et al., 2002; Shan, 2016; Wennerberg et al., 2005). Recently, however, DNA partitioning proteins of the ParB family have emerged as a novel class of molecular switches that require the pyrimidine nucleotide cytidine triphosphate (CTP) for proper function (Osorio-Valeriano et al., 2019; Soh et al., 2019).

ParB forms part of the ParAB*S* system, which is highly conserved among bacteria and serves to actively segregate newly replicated chromosomes and low-copy number plasmids to the nascent daughter cells (Badrinarayanan et al., 2015). It is a dimeric DNA-binding protein that recognizes clusters of short palindromic sequences (*parS*) close to the replication origin of target DNA molecules (Lin and Grossman, 1998; Mohl and Gober, 1997). After initial specific interaction with individual *parS* sites, ParB spreads into adjacent DNA segments, giving rise to a large nucleoprotein assembly (partition complex) that typically includes 10-20 kb of the origin region (Breier and Grossman, 2007; Lynch and Wang, 1995). Upon DNA replication, the partition complexes on sister DNA molecules interact dynamically with the P-loop ATPase ParA, thereby promoting the movement of the associated origin region towards opposite cell poles. The segregation process is driven by a ratchet-like mechanism, in which ParB follows a retracting polarized gradient of nucleoid-associated ParA molecules (Fogel and Waldor, 2006; Lim et al., 2014; Vecchiarelli et al., 2014; reviewed by Jalal and Le, 2020). In addition to its role in origin segregation, the partition complex can have several additional functions (Kawalek et al., 2020). In many bacteria, it interacts with polar scaffold proteins to anchor the origin regions at the cell poles once their segregation is finished (Bowman et al., 2008; Donovan et al., 2012; Ebersbach et al., 2008; Yamaichi et al., 2012). Moreover, it often serves as a loading platform for the SMC condensin complex, which facilitates bulk chromosome segregation by aligning the two arms of newly replicated sister chromatids (Böhm et al., 2020; Gruber and Errington, 2009; Sullivan et al., 2009; Tran et al., 2017). Finally, in certain species, ParB acts as a polar landmark that drives the establishment of a regulatory protein gradient controlling division site placement (Thanbichler and Shapiro, 2006; Toro-Nahuelpan et al., 2019).

The precise architecture of partition complexes has long been a matter of debate (Jalal and Le, 2020). In particular, it has remained unclear how a small number of *parS* sites, or even a single copy (Graham et al., 2014), can be sufficient to recruit several hundred ParB molecules to the origin region. Several models have been proposed to explain the underlying mechanism, including the polymerization of ParB at *parS* sites (Murray et al., 2006; Rodionov et al., 1999), bridging interactions between distal ParB proteins (Broedersz et al., 2014; Debaugny et al., 2018; Graham et al., 2014; Sanchez et al., 2013; Song et al., 2017; Taylor et al., 2015) and the assembly of ParB into biomolecular condensates (Guilhas et al., 2020). Fundamental new insights into the underlying mechanism came from recent work demonstrating that ParB is a CTPase that acts as a DNA-sliding clamp (Jalal et al., 2020a; Osorio-Valeriano et al., 2019; Soh et al., 2019). This property is closely linked to its highly conserved domain structure (Funnell, 2016) (**Figure 1A**), which includes (1) an N-terminal nucleotide-binding (NB) domain (Osorio-Valeriano et al., 2019; Soh et al., 2019), (2) a central helix-turn-helix (HTH) DNA-binding domain mediating *parS* recognition (Chen et al., 2015; Leonard et al., 2004; Schumacher et al., 2010; Surtees and Funnell, 2001), (3) a non-structured linker region and (4) a C-terminal dimerization domain that tightly links the two subunits of a ParB dimer (Fisher et al., 2017; Leonard et al., 2004). It has been shown that, once the DNA-binding domain of ParB interacts with *parS*, the two NB domains associate with each other in a CTP-dependent manner, thereby closing the ParB dimer at both ends, with DNA entrapped in between the two subunits (Osorio-Valeriano et al., 2019; Soh et al., 2019). ParB rings have a reduced affinity for *parS*. As a consequence, they leave their initial binding site and slide into the flanking DNA regions, making *parS* again accessible to other ParB dimers (de Asis Balaguer et al., 2021; Jalal et al., 2020a; Soh et al., 2019). In this way, a single *parS* motif can act as a loading site for many ParB rings.

**Figure 1.**
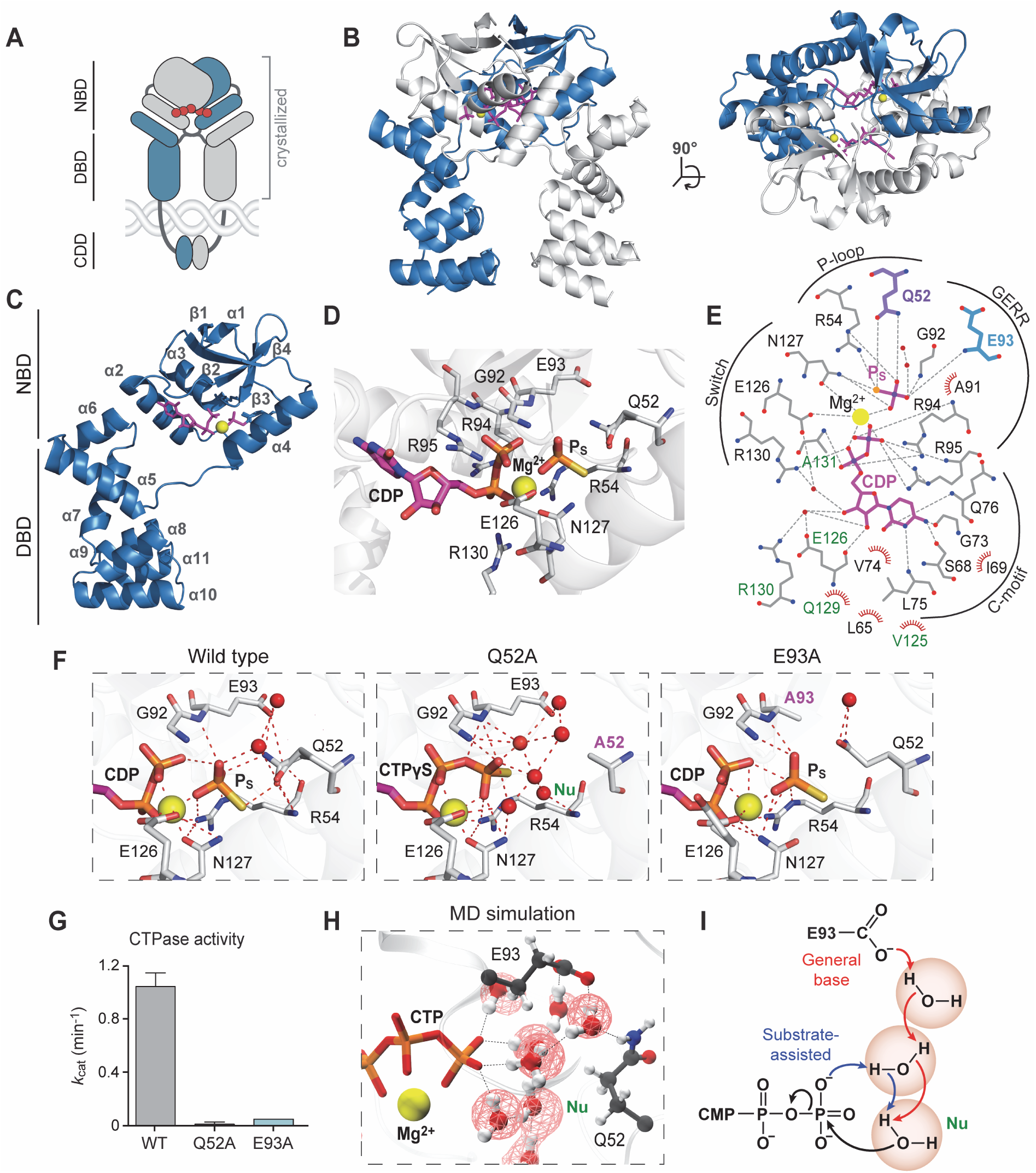
Crystal structures of *Mx*ParB in the pre- and post-hydrolysis state provide insight into the mechanism of CTP hydrolysis. **(A)** Scheme of a closed ParB clamp entrapping DNA. The two subunits are colored blue and gray, respectively. CTP molecules are depicted as red filled circles. The regions corresponding to the nucleotide-binding domain (NBD), DNA-binding domain (DBD) and C-terminal dimerization domain (CDD) are indicated. **(B)** Side view (left) and top view (right) of the crystal structure of dimeric nucleotide-bound ParB (NBD and DBD) in ribbon representation. One subunit is colored in blue and the second subunit in gray. CDP and monothiophosphate (P_S_) are depicted in magenta, and Mg^2+^ ions are shown in yellow. **(C)** Arrangement of secondary structure elements in a single ParB subunit. **(D)** Magnification of the nucleotide-binding pocket of ParB. Residues interacting with the diphosphate moiety of CDP and P_S_ are shown in stick representation. The Mg^2+^ ion is colored yellow. **(E)** Protein-ligand interaction 2D map of CDP and P_S_ bound to ParB. Hydrogen bonds (shorter than 3.02 Å) are shown as dashed gray lines and hydrophobic interactions as red semicircles; water molecules are depicted as red solid circles. Nitrogen and oxygen atoms are indicated in blue and red, respectively. Residues from the *cis-* subunit are labeled in black, those from the *trans-*subunit in green. The catalytic residues Q52 and E93 are highlighted in purple and blue, respectively. Conserved motifs are indicated. **(F)** Magnification of the catalytic site of wild-type *Mx*ParB and its catalytically inactive Q52A and E93A variants. Residues interacting with the Mg^2+^ ion and P_S_ (wild-type and E93A) or the γ-phosphate moiety of CTP (Q52A) are shown as sticks. Water molecules are depicted as red spheres. The putative nucleophilic water (Nu) in the Q52A structure is labeled in green. Mutated residues are labeled in magenta. **(G)** CTPase activities of mutant ParB variants. The indicated proteins (5 μM) were incubated with 1 mM CTP in the presence of *parS* DNA (250 nM). The reaction rates were determined with a NADH-coupled enzyme assay. Data represent the mean of three replicates (± SD). **(H)** Snapshot from MD simulations showing the catalytic site of wild-type ParB in the pre-hydrolysis state. For clarity, only the triphosphate moiety, the Mg^2+^ ion, the water molecules and the catalytic residues Q52 and E93 are shown. Water occupancies are depicted as red wire frames. The putative nucleophilic water is labeled in green (Nu). Hydrogen bonds are indicated by red dotted lines. **(I)** Proposed catalytic mechanism of CTP hydrolysis by ParB-type CTPases. E93 acts as a general base that deprotonates the nucleophile through a water wire. In its absence, an alternative, substrate-assisted mechanism could be responsible for the residual hydrolytic activity observed for the E93A variant. See also Figures S1 and S2.

While the function of ParB as a molecular sliding-clamp is well-established, the molecular mechanisms controlling its loading, movement and release are still incompletely understood. In this study, we use a broad, interdisciplinary approach to dissect the CTPase cycle of ParB and its physiological significance in the model organism *Myxococcus xanthus*. We solve crystal structures of ParB in the pre- and post-hydrolysis state and provide insight into the catalytic mechanism of CTP hydrolysis. Moreover, we investigate the effects of *parS* and CTP binding on the conformational dynamics of ParB in solution and thus shed light on the mechanism of *parS*-triggered ring closure. Based on the structural data, we generate ParB variants that lack CTPase activity and are thus locked in the closed (ring) state. Using these proteins, we show that CTP hydrolysis is important, albeit not essential for the dissociation of ParB from DNA *in vitro*. Consistent with this finding, its loss leads to a more stable association and wider spreading of ParB within the chromosomal origin region *in vivo*, pointing to key role of CTP hydrolysis in ParB ring opening and release. Notably, although CTPase-deficient ParB variants still form well-defined (albeit larger) partition complexes within the cell, they are no longer able to support chromosome segregation, with their expression leading to cell death. Collectively, our results provide comprehensive new insights into the structural basis of ParB function and the role of its CTPase cycle in partition complex dynamics, thereby furthering our understanding of this novel molecular switch.

## RESULTS

### X-ray crystallography reveals the structure of *Mx*ParB in the post-hydrolysis state

Recent work solved the crystal structure of a truncated variant (amino acids 21-208) of ParB from *B. subtilis* (*Bs*ParB) in the closed (ring) conformation (Soh et al., 2019). It revealed that the interaction between the two NB domains is mediated by two nucleotide molecules that are sandwiched at the subunit interface. However, the nucleotide detected was cytidine diphosphate (CDP), although CTP was added to the crystallization buffer, suggesting that CTP was hydrolyzed after crystal growth. A second study solved the structure of a homologous NB domain from PadC (Osorio-Valeriano et al., 2019), an accessory protein of the ParAB*S* system in *M. xanthus* (Lin et al., 2017). Although this structure contains CTP and is highly similar to that of the corresponding domain from ParB, the environment of the triphosphate moiety shows significant differences, in line with the fact that the NB domain of PadC lacks CTPase activity and forms a constitutive dimer. Thus, collectively, the precise nature of the ParB CTP-binding pocket, and in particular the catalytic region surrounding the γ-phosphate moiety, remained unclear.

To address this issue, we purified a truncated variant of *M. xanthus* ParB comprising only the NB and DNA-binding domains (amino acids 35-246) and crystallized it in the presence of the poorly hydrolyzable CTP analog CTPγS (Osorio-Valeriano et al., 2019). The resulting X-ray structure (1.9 Å resolution) is similar to that of the *Bs*ParB·CDP complex (RMSD 0.889 Å for 194 paired C_α_ atoms) and shows a dimer composed of two tightly interlinked NB domains followed by adjacent, spatially separated DNA-binding domains (**Figures 1B** and **S1**; **Table S1**). The NB domain contains a conserved ParB/sulfiredoxin (ParB/Srx) fold composed of a four-stranded β sheet, in which strand β1 is connected to β2 by a large loop containing helices α1 and α2 and strands β3 and β4 are connected by helix α3 (**Figure 1C**). This module is connected to helix α4, which lines the surface of strands β2 and β3 and closes off the nucleotide-binding site. A long non-structured region between helices α4 and α5 then links the NB domain to the DNA-binding domain, which is composed of seven densely packed helices (α5-α11), with helices α6 and α7 forming the characteristic HTH motif. In the dimeric complex, the two polypeptide chains cross over at this linker region, so that helix α4 of each chain is placed next to helix α5 of the *trans*-subunit. As a consequence, the two NB domains adopt a face-to-face arrangement, with the two nucleotides enclosed adjacent to each other at the domain interface (**Figures 1B** and **1C**). The high resolution of our structure allowed us to clearly identify the ligands and water molecules in the nucleotide-binding pocket. Surprisingly, we did not detect CTPγS but CDP, indicating that the nucleotide had again been hydrolyzed during crystallization (**Figures 1D-1F** and **S2A**). However, unlike in the *Bs*ParB·CDP structure (Soh et al., 2019), the resulting thiophosphate groups were still associated with the protein, thus providing insight into the post-hydrolysis state of *Mx*ParB. The nucleotides, complexed with Mg^2+^ ions, are located in pockets that are formed by helices α2-α4 and loop α2/β2 of one subunit and helix α4 of the respective *trans*-subunit. They form a dense network of hydrogen bonds with amino acid residues and backbone groups in these regions, thereby tightly linking the two adjacent subunits. The nucleobase occupies a cavity that is just large enough to fit a pyrimidine ring, thereby excluding the larger purine nucleotides ATP and GTP. Specificity for cytosine nucleotides is likely provided by hydrogen-bonding interactions of the 4-amino group of the cytosine moiety with the hydroxy group of S68 and the backbone carbonyl group of G73. Overall, the architecture of the nucleotide-binding site is very similar to that of *Bs*ParB (**Figure S1**), indicating that the mode of nucleotide binding is highly conserved among members of the ParB family. Notably, the thiophosphate generated during CTPγS hydrolysis remained bound to the Mg^2+^ ion and, thus, in close vicinity of the diphosphate moiety of CDP (**Figures 1D-1F**). It is held in place by hydrogen bonds with the backbone amides of G92 and E93 at the beginning of helix α3, with residue N127 in helix α4 and with residues Q52 and R54 in the large loop α1/α2, which forms a lid-like structure covering the active site. In the absence of crystal packing constraints, these interactions may promote conformational changes in these regions that promote the release of the leaving (thio)phosphate and thus stimulate ring opening. Notably, all residues that are in contact with the thiophosphate molecule are highly conserved in ParB proteins (**Figure S1A**), suggesting that they have a critical role in their function.

### CTP hydrolysis involves a water network connected to the catalytic residues Q52 and E93

The thiophospate-binding residues Q52 and E93 have not been previously implicated in ParB function. Their side chains are too remote to interact directly with the bound nucleotide, but their proximity to the leaving nucleotide γ-phosphate opens the possibility that they could have a role in the mechanism of nucleotide hydrolysis. To test this hypothesis, we generated *Mx*ParB variants with alanine substitutions at these positions (Q52A or E93A) and determined their ability to hydrolyze CTP in reactions containing *parS* to stimulate the homodimerization of the NB domains. Notably, both variants showed a dramatic decrease in the CTP turnover rate (**Figure 1G**), although they still bound nucleotides with approximately wild-type affinity (**Figure S2B**), suggesting a defect in their catalytic activity. Prompted by these results, we extended our structural analyses to the two mutant proteins. For this purpose, we generated truncated derivatives comprising only the NB and DNA-binding domains, crystallized them in the presence of CTPγS and solved their X-ray structures to 1.5 Å (Q52A) and 1.9 Å (E93A) resolution (**Table S1**). Both proteins formed dimeric complexes highly similar to the one obtained for the wild-type protein, with RMSD values of 0.54 Å (Q52A; for 194 paired C_α_ atoms) and 0.59 Å (E93A; for 193 paired C_α_ atoms), respectively. As expected, the Q52A variant contained intact CTPγS and thus appeared to be locked in the pre-hydrolysis state (**Figures 1F** and **S2A**). By contrast, the E93A variant was again associated with CDP and thiophosphate, suggesting that its slightly higher residual CTPase activity (see **Figure 1G**) was sufficient to achieve nucleotide hydrolysis during crystallization. In both cases, the architecture of the nucleotide-binding pocket was largely unchanged compared to the wild-type *Mx*ParB·CDP·P_S_ complex. The only exception is an outward rotation of residue Q52 in the E93A variant, likely caused by the loss of the hydrogen bond formed between Q52 and E93 in the wild-type protein (**Figure S2C**). The mutation of E93 also changed the orientation of the adjacent residue R97, which normally forms a π-stacking interaction with residue F57 in loop α1/α2 that may be required to stabilize the lid covering the active site (**Figure S2C**). In the *Mx*ParB_Q52A_·CTPγS structure, residues R54, G92, E93 and N127, which bind the thiophosphate molecule in the post-hydrolysis complexes, interact with the γ-thiophosphate group of CTPγS (**Figures 1E-1F**). Notably, in the absence of a catalytic product, the cavity between the nucleotide and loop α1/α2 is occupied by a network of six water molecules that form hydrogen bonds with the γ-thiophosphate, the main chain of residues R54 and I90 as well as the side chains of E126, N127 and, importantly, E93 (**Figure 1F**). One of these waters is positioned in line with the scissile bond, at a position that would allow a nucleophilic attack on the γ-(thio)phosphate group. These findings suggest that residues Q52 and E93 may mediate the activation of the nucleophile by affecting the properties of this water network.

To more closely investigate the possible role of the water network and the surrounding active site residues in the mechanism of nucleotide hydrolysis, we performed molecular dynamics (MD) simulations of *Mx*ParB in the CTP-bound, pre-hydrolysis state. The MD simulations were initiated from the wild-type and mutant protein structures obtained in this study (**Figure 1**), with CTPγS or CDP·P_S_ converted to CTP based on the arrangement of the nucleotide in the *Mx*ParB_Q52A_·CTPγS complex. The results showed that the network of six water molecules identified in the Q52A structure was retained under physiological conditions in both the wild-type protein (**Figure 1H**) and the Q52A and E93A variants (**Figure S2D**). Although the water network was dynamic, as expected due to the thermal fluctuations at room temperature, the average positions of the water molecules in the simulations agreed with the positions of the crystal water molecules, as shown by the well-defined spatial densities in the corresponding occupancy maps (**Figures 1H** and **S2D**). This is also true for the putative nucleophilic water, which was connected to residue E93 through a water wire. The simulations also identified a seventh water molecule close to the carboxyl group of residue E93, albeit with lower occupancy. We speculate that E93 could act as a catalytic base that abstracts a proton from an adjacent water and, transmitted through a proton wire, promotes the nucleophilic attack of a polarized water molecule on the γ-phosphate group (**Figure 1I**). Interestingly, although the E93A variant shows a drastically reduced turnover rate, it retains some residual activity (**Figure 1G**). This observation may be explained by the existence of an alternative, albeit less efficient, substrate-assisted hydrolysis pathway, in which the γ-phosphate group acts as a catalytic base (Hanekop et al., 2006; Pasqualato and Cherfils, 2005; Schweins et al., 1995; Schweins et al., 1994; Zaitseva et al., 2005) (**Figure 1I**). The role of residue Q52 is less clear. In the *Mx*ParB·CDP·P_S_ structure, it interacts tightly with the leaving thiophosphate molecule (**Figures 1E** and **1F**), suggesting that it may be required to bind and stabilize the catalytic product and, thus, slow down the reverse reaction. Notably, the MD simulations show that, in the Q52A variant, the lid structure formed by loop α1/α2 (residues 56-61) is significantly more rigid than in the wild-type protein (**Figure S2E**). The absence of Q52 may thus abolish nucleotide hydrolysis both by preventing the release of the leaving phosphate and promoting its transfer back to the CDP molecule. Apart from that, in the pre-hydrolysis complex, Q52 can form a hydrogen bond with residue E93 (**Figures 1H** and **S2C**) and may keep its carboxyl group in an orientation optimal for its interaction with the water network. Interestingly, in the MD simulations, the side chain of residue Q52 is flexible and alternates between configurations in which it points toward or away from the γ-phosphate group and the associated water network (**Figure S2E**). Thus, it is conceivable that configurations that favor the hydrolytic reaction are adopted only rarely, contributing to the low catalytic rate of *Mx*ParB. Furthermore, consistent with the crystallographic data (**Figure S2C**), the simulations show that the loss of E93 increases the flexibility of residues Q52 and R97 and thus destabilizes loop α1/α2 in the region surrounding F57 (**Figure S2E**). Collectively, these results provide first insights into the possible catalytic mechanism of ParB proteins and thus set the basis for in-depth experimental and quantum mechanics/molecular mechanics (QM/MM) studies of nucleotide hydrolysis by this new family of NTPases.

### CTP and *parS* binding promote ring formation by inducing synergistic conformational changes

The nucleotide-bound, closed state of ParB is characterized by a loose connection between the NB and DNA-binding domains in *cis*, enabling ring closure through interactions of helix α4 with helices α5’ and α6’ of the *trans*-subunit (**Figure 1B**; (Soh et al., 2019)). Interestingly, however, in the previously reported apo structures of *Bs*ParB (Leonard et al., 2004) and its close homolog Noc (Jalal et al., 2021), the two domains are rotated by 103° relative to each other, forming a compact unit in which helix α4 bundles with helices α5 and α6 of the same polypeptide chain (**Figure S3A**; see **Figure 2A** for a homology model of the corresponding *Mx*ParB structure). This observation suggests that the transition of ParB from the open to the closed state may involve a conformational switch in its subunits, likely driven by CTP and/or *parS* binding. To investigate this possibility, we analyzed the behavior of *Mx*ParB at different stages of the DNA loading process using hydrogen-deuterium exchange (HDX) mass spectrometry (**Figures 2B-2D**, **S3** and **S4**). This technique detects local shifts in the accessibility of backbone amide hydrogens caused by conformational changes or ligand binding and, thus, provides a powerful means to resolve the structural dynamics of proteins in solution (Konermann et al., 2011).

**Figure 2.**
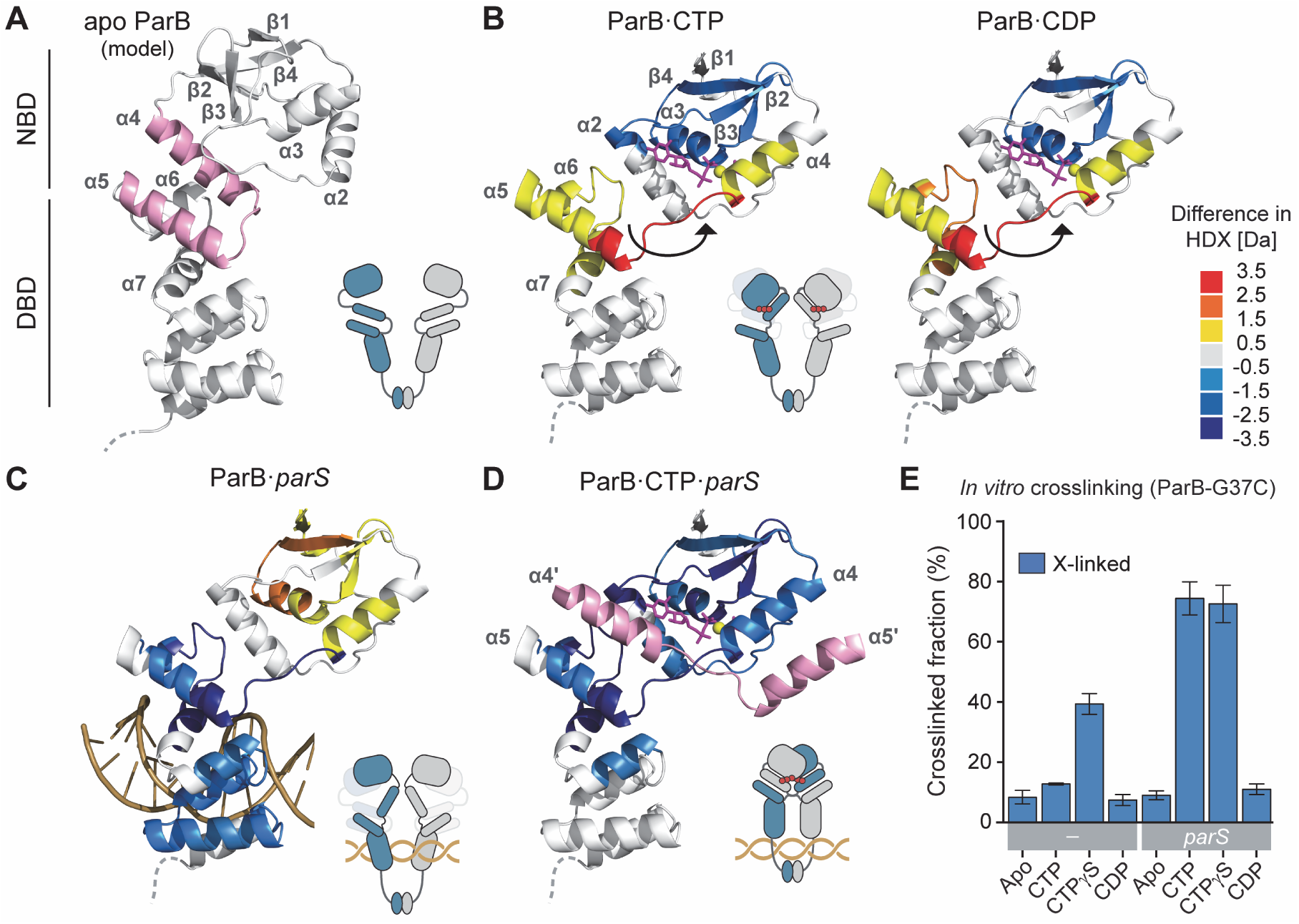
CTP and *parS* induce synergistic conformational changes required for ParB clamp closure. **(A)** Model of apo *Mx*ParB (residues V38 to Q229) based on the apo structure of the ParB homolog Noc from *Geobacillus thermoleovorans* (PDB: 7NFU). Helices α4 and α5 are highlighted in pink. Regions corresponding to the N-terminal NB domain (NBD) and the DNA-binding domain (DBD) are indicated. **(B)** Change in hydrogen-deuterium exchange (HDX) upon incubation of *Mx*ParB with CTP or CDP, mapped onto the crystal structure of nucleotide-bound ParB (t = 30 s; **Figure S4**) for better visualization. Only one subunit is shown for simplicity. Black arrows indicate the release of the NB domain. **(C)** Change in HDX upon incubation of *Mx*ParB with a *parS-*containing oligonucleotide (t = 30 s; **Figure S4)**. A DNA molecule was modeled into the structure, based on the crystal structure of the *H. pylori* ParB·*parS* complex (PDB: 4UMK). **(D)** Differences in HDX upon incubation of ParB with a *parS* DNA in the presence of CTP (t = 30 s; **Figure S4)**. Helices α4 and α5 from the second subunit are shown in pink. The color code for all HDX analyses is given on the top right. Cartoons represent hypothetical conformations of the ParB dimer inferred from the HDX data. **(E)** Efficiency of *Mx*ParB ring closure in different conditions. ParB-G37C was incubated with the indicated nucleotide (1 mM) in the absence or presence of *parS* DNA prior to crosslinking with BMOE. After separation of the reaction products by SDS-PAGE (**Figure S3B**), the fraction of crosslinked (X-linked) protein obtained in each condition was determined by densitometric analysis. Data represent the mean of three replicates (± SD). See also **Figures S3-S5**.

First, we compared the HDX profiles of *Mx*ParB in the apo and nucleotide-bound state (**Figures 2A** and **2B**). As expected, the addition of CTP led to the protection of regions lining the nucleotide-binding site, in particular helix α3 and sheets β2-β4. However, importantly, we also observed a strong deprotection of helices α4-α6 at the junction of the NB and DNA-binding domains. This observation suggests that CTP binding indeed promotes the dissociation of the two domains, thereby exposing helices α4 and α5/α6 to the environment and increasing their HDX rates. The driving force behind this conformational change may be interactions of the CTP triphosphate moiety and the Mg^2+^ ion with residues E126, N127 and R130 at the end of helix α4 (**Figures 1C-1E**) that destabilize the α4-α5/α6 interface. To correlate these results with the opening state of *Mx*ParB, we additionally performed site-specific chemical crosslinking studies. For this purpose, we introduced a cysteine residue (G37C) at the N-terminal end of the NB domain, close to the symmetry axis of the closed complex. Upon ring closure, the corresponding residues of the two subunits are placed next to each other at a distance that enables their covalent crosslinking by the bifunctional thiol-reactive compound bismaleimidoethane (BMOE). This approach confirmed that the CTP-induced release of the NB domain alone is not sufficient to induce ring closure (**Figures 2E and S3B**). Notably, CDP produces a similar change in the HDX profile, although it causes less protection of the NB domain (**Figure 2B**), likely due to its considerably lower binding affinity (Osorio-Valeriano et al., 2019). Very similar results were obtained with a truncated *Mx*ParB variant lacking the C-terminal dimerization domain (*Mx*ParB_ΔC_), which verifies that the effects observed were independent of intra-subunit interactions (**Figure S5**).

Next, we reinvestigated the structural consequences of *parS* binding (**Figure 2C**). Consistent with our previous results (Osorio-Valeriano et al., 2019), we detected a strong protection of the DNA-binding domain, especially in regions previously shown to interact directly with the associated DNA molecule (Chen et al., 2015; Jalal et al., 2020b; Sanchez et al., 2013). In addition, we also observed the deprotection of regions in the NB domain, including helices α3 and α4 and sheets β1-β4. This result may again indicate the release of helix α4 from the DNA-binding domain, triggered by *parS*-induced conformational changes in helices α5 and α6. Notably, in crystal structures of ParB·*parS* complexes (Chen et al., 2015; Jalal et al., 2020b), helices α4-α6 still adopt an arrangement similar to that in the apo state (**Figure S3C**). We hypothesize that *parS* binding weakens the interactions in this region but does not fully abolish them. In solution, the NB domain may thus alternate between the tethered and released state, although crystal packing may favor the more compact tethered state. The transition to the released state provides the basis for face-to-face interactions between adjacent NB domains. This process may be facilitated by accompanying structural changes that modulate the architecture of the nucleotide-binding pocket or fine-tune the arrangement of helices α5 and α6 such as to promote their interaction with helix α4 of the *trans-*subunit. The concomitant binding of CTP may fully stabilize the released state. Moreover, it is required to complete the dimerization surface, thereby finally enabling ring closure (**Figures 2E** and **S3B**).

The synergistic effect of *parS* and CTP on ring closure was evident from the HDX profile of *Mx*ParB in reactions containing both ligands (**Figure 2D**). In this condition, the entire NB domain was strongly protected, consistent with a close association of the two subunits in this region. Protection was also observed in the helix α5/α6 region of the DNA-binding domain, reflecting its interaction with the swapped helix α4, as well as in segments of helices α7 and α8 that are in close vicinity of helix α5. Notably, identical HDX profiles were obtained when *Mx*ParB was incubated with both *parS* and CTPγS or even with CTPγS alone (**Figures S3D and S3E**), consistent with the strong stimulatory effect of this slowly hydrolyzable nucleotide analog on ring closure (Jalal et al., 2020a; Osorio-Valeriano et al., 2019; Soh et al., 2019) (**Figures 2E** and **3A**). CDP, by contrast, did not promote the transition to the closed state when combined with *parS* (**Figure S3F**). HDX analysis of a truncated *Mx*ParB variant (*Mx*ParB_ΔC_) showed that CTPγS was not able to promote self-dimerization of the NB domains in the absence of a C-terminal linkage and rather produced a protection pattern comparable to that of CTP (**Figure S5**). The *parS*-independent self-association of the NB domains observed in the absence of nucleotide hydrolysis is thus aided by the proximity of the two subunits mediated by the C-terminal dimerization domain. Interestingly, in the presence of *parS*, the closed complexes of *Mx*ParB (**Figures 2D** and **S3E**) did not display the typical global protection of the DNA-binding domain observed for the open dimers (**Figures 2C** and **S3F**). This result supports the previous finding that ring closure reduces the affinity of ParB for *parS* sites (Jalal et al., 2020a; Osorio-Valeriano et al., 2019; Soh et al., 2019). Thus, ParB loading is a multi-step process that is driven by a complex and synergistic interplay of the two subunits with each other and their ligands CTP and *parS*.

**Figure 3.**
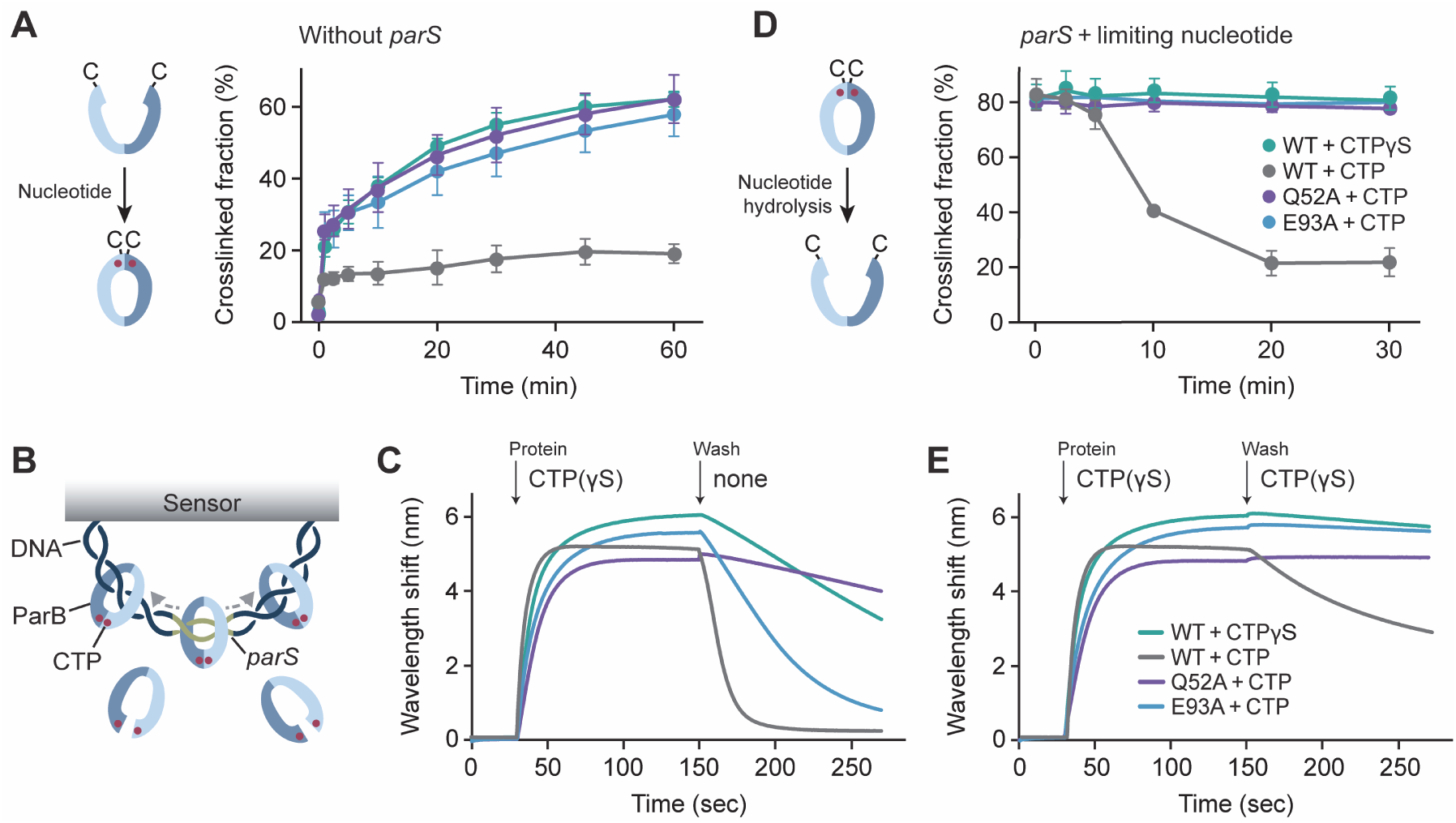
CTP hydrolysis triggers the opening of ParB rings and their unloading from DNA *in vitro*. **(A)** Site-specific crosslinking analysis of ParB variants harboring an additional G37C substitution in the absence of *parS* DNA. Mutant ParB variants were incubated with 1 mM CTP or CTPγS. Aliquots were withdrawn at the indicated time points, treated with the thio-reactive crosslinker BMOE and analyzed by SDS-PAGE. The graph shows the fraction of crosslinked protein in each of the samples. Data represent the mean of three replicates (± SD). The legend is shown in Figure 3D. **(B)** Scheme of the biolayer interferometry (BLI) setup used to analyze the CTP-dependent accumulation of *Mx*ParB on closed DNA substrates. Double-biotinylated dsDNA fragments (234 bp) containing a single *parS* site were immobilized on streptavidin-coated BLI sensors to generate closed substrates. The sensors were then probed with *Mx*ParB to analyze its nucleotide-dependent loading and dissociation. **(C)** BLI analysis of the interaction of *Mx*ParB with closed DNA substrates. Sensors carrying closed *parS*-containing dsDNA fragments were probed with the indicated ParB variants in the presence of 1 mM CTP or CTPγS. At the end of the binding reaction, the sensors were transferred into protein- and nucleotide-free buffer to follow the dissociation reactions. The kinetics of the loading and unloading processes were followed by monitoring the wavelength shifts induced by changes in the optical thickness of the sensor surface during the association and dissociation phases. The legend is shown in (E). **(D)** Site-specific crosslinking analysis of CTPase-dependent *Mx*ParB ring opening. The indicated ParB variants (containing an additional G37C mutation) were pre-incubated for 5 min with low concentrations of CTP or CTPγS (100 µM) in the presence of *parS* DNA (1 µM) to induce the formation of ParB rings. Subsequently, samples were taken at the indicated time points, treated with BMOE and analyzed as described in (A). Data represent the mean of three replicates (± SD). Note that *Mx*ParB rings open gradually as CTP is consumed. **(E)** BLI analysis of the dissociation of *Mx*ParB rings from DNA in the presence of nucleotide. The analysis was performed as described in Figure 3C, with the exception that 1 mM nucleotide was included in the dissociation buffer. See also **Figures S6**.

### CTP hydrolysis triggers opening of ParB rings and their release from DNA

Our analyses revealed Q52 and E93 as catalytic residues. Their exchange largely abolished nucleotide hydrolysis but still allowed homodimerization of the NB domains during crystallization. To analyze the conformational dynamics of the Q52A and E93A variants in solution, we introduced a cysteine residue in their N-terminal region (G37C) and probed their opening state by BMOE-mediated chemical crosslinking. Both mutant variants (Q52A, E93A) were able to form rings in the presence of CTP, even if *parS* was omitted from the reactions (**Figures 3A and S6A**). Wild-type *Mx*ParB (G37C), by contrast, only showed marginal crosslinking under this condition, consistent with its reported dependency on *parS* for ring closure (Osorio-Valeriano et al., 2019). However, it displayed the same behavior as the two CTPase-deficient variants when incubated with the poorly hydrolyzable nucleotide analog CTPγS. These findings verify the ability of the Q52A and E93A variants to transition to the closed state in solution, and they suggest a critical role of nucleotide hydrolysis in the dynamics of ring formation. The identification of the Q52A and E93A variants enabled us to investigate the role of nucleotide hydrolysis on the DNA-loading process. For this purpose, we adapted a previously published biolayer interferometry (BLI) assay (Jalal et al., 2020a), monitoring the accumulation of ParB rings on a closed *parS*-containing DNA fragment that was immobilized on a sensor surface (**Figure 3B**). As reported for *Cc*ParB (Jalal et al., 2020a), wild-type *Mx*ParB was efficiently loaded onto DNA in the presence of CTP or CTPγS. By contrast, only a weak signal was observed without nucleotide, likely reflecting the sequence-specific binding of a single, open dimer to *parS* (**Figure S6C**). Notably, the Q52A and E93A variants were still loaded with wild-type efficiency, confirming that the mutations did not impair ring closure but specifically affected the catalytic activity of *Mx*ParB. We then followed the dissociation of proteins from loaded DNA fragments after transfer of the BLI sensor into protein- and nucleotide-free buffer (**Figure 3C**). Whereas wild-type *Mx*ParB rings assembled in the presence of CTP were released at an appreciable rate (*k_off_* = 0.12 s^-1^), dissociation slowed down considerably (∼22-fold) when the protein was loaded with the slowly hydrolyzable nucleotide analog CTPγS (*k_off_* = 0.005 s^-1^). Importantly, a severe reduction in the dissociation rate was also observed for the catalytically impaired Q52A (*k_off_* = 0.002 s^-1^) and, to a lesser extent, E93A (*k_off_* = 0.018 s^-1^) variants, even when loaded in the presence of CTP. Thus, nucleotide hydrolysis is required to efficiently open *Mx*ParB rings and release them from DNA. Consistent with this result, *in vitro* crosslinking studies showed that *Mx*ParB rings formed in the presence of CTP or CTPγS and a short, *parS*-containing DNA stemloop only returned back to the open state under conditions that allowed hydrolysis of the bound nucleotide (**Figure 3D**).

Interestingly, in the BLI experiments, ring release was strongly influenced by the presence of nucleotide during the dissociation phase. In nucleotide-free buffer, all locked proteins still dissociated to an appreciable extent, despite the lack of CTPase activity (**Figure 3C**). In the presence of saturating nucleotide concentrations, by contrast, they remained stably associated with the DNA, and ring release was strictly dependent on nucleotide hydrolysis (**Figure 3E**). Titration experiments showed that the rate of ring release increased gradually with decreasing nucleotide concentrations in the wash buffer (**Figure S6B**). These results suggest that *Mx*ParB rings can open either through spontaneous dissociation of the NB domains, followed by the loss of the bound nucleotide, or through nucleotide hydrolysis. In nucleotide-containing buffer, spontaneous nucleotide loss may be minimized, enabling the NB domains to reassociate before ParB is released from the DNA. Nucleotide hydrolysis, by contrast, may trigger a conformational change that induces the transition of ParB to a permanently open state, necessitating its interaction with *parS* for reloading.

To further probe the mechanism of *Mx*ParB function, we extended the BLI analysis to a series of additional *Mx*ParB variants (R54A, R94A, R95A, E126A, N127A, R130A) with mutations at positions close to the γ-phosphate moiety of the bound nucleotide. However, all of these proteins were unable to accumulate on DNA (**Figure S6C**), because they either were defective in nucleotide binding (R95A, R130A) (Osorio-Valeriano et al., 2019) or still bound nucleotide but failed to undergo ring closure (R54A, R94A, E126A, N127A), as suggested by their lack of CTPase activity (**Figures S6D** and **S6E**; **Table S2**). Thus, the Q52A and E93A exchanges are the only mutations that lock *Mx*ParB in the nucleotide-bound, closed state.

Notably, an *Mx*ParB variant lacking the dimerization domain (*Mx*ParB_ΔC_) was no longer able to accumulate on a closed *parS*-containing DNA fragment (**Figure S6C**), even though it bound nucleotide with wild-type affinity (**Figure S6F**) and still showed *parS*-dependent CTPase activity (**Figure S6G**), indicating that the juxtaposition of two ParB monomers at the palindromic *parS* site was sufficient to promote the dimerization of their NB domains. We further observed that the C-terminal dimerization domains of a full-length *Mx*ParB derivative carrying an engineered C-terminal cysteine residue (G296C) were crosslinked with high efficiency, even in the absence of CTP (**Figure S6H**), in line with the notion that they are stably bound to each other and seal the ParB ring tightly at the C-terminal end (Fisher et al., 2017; Leonard et al., 2004; Taylor et al., 2021). Together, these results confirm that the C-terminal regions form an essential part of the ring structure that embraces DNA after the release of ParB from the *parS* binding site. Nucleotide hydrolysis then acts as a switch to open the N-terminal gate, thereby allowing the release of the bound DNA and the dissociation of ParB from the partition complex.

### ParB CTPase activity controls the size of the partition complex

Our *in vitro* studies showed that CTP and *parS* binding act synergistically to drive *Mx*ParB into the closed-ring state, whereas CTP hydrolysis promotes ring opening and release. To clarify the physiological relevance of this nucleotide-regulated conformational cycle, we next investigated the ability of CTPase-deficient *Mx*ParB variants to form partition complexes *in vivo*. To this end, wild-type *Mx*ParB and various mutant variants were N-terminally fused to a hemagglutinin (HA) affinity tag and produced in cells depleted of the native ParB protein (**Figure S7A**). Subsequently, the distribution of the different fusion proteins within the chromosomal origin region was determined by chromatin immunoprecipitation followed by high-throughput sequencing of the isolated DNA fragments (ChIP-seq). The *M. xanthus* chromosome contains a total of 24 *parS* sites, most of which are densely clustered within a 3 kb region at a distance of ∼30 kb from *oriC* (Harms et al., 2013; Iniesta, 2014). Consistent with results obtained in other species (Böhm et al., 2020; Breier and Grossman, 2007; Tran et al., 2018), the tagged wild-type protein formed a compact peak with a width of 33 kb that was centered at the *parS* cluster (**Figure 4A**). The catalytically inactive Q52A variant, by contrast, displayed a considerably broader distribution and covered up to 175 kb of the origin region. A similar behavior, albeit with a more narrow distribution (125 kb), was observed for the E93A variant, whereas an *Mx*ParB variant defective in ring closure due to a defect in CTP binding (R95A; see **Figure S6C** and **Table S2**) (Osorio-Valeriano et al., 2019) did not show any accumulation in the origin region (**Figure S7B**). Notably, the enhanced spreading of the Q52A and E93A variants correlated with their slower release from DNA *in vitro* (compare **Figures 3C** and **3E**), indicating a close link between the stability of ParB rings and the size of partition complexes.

**Figure 4.**
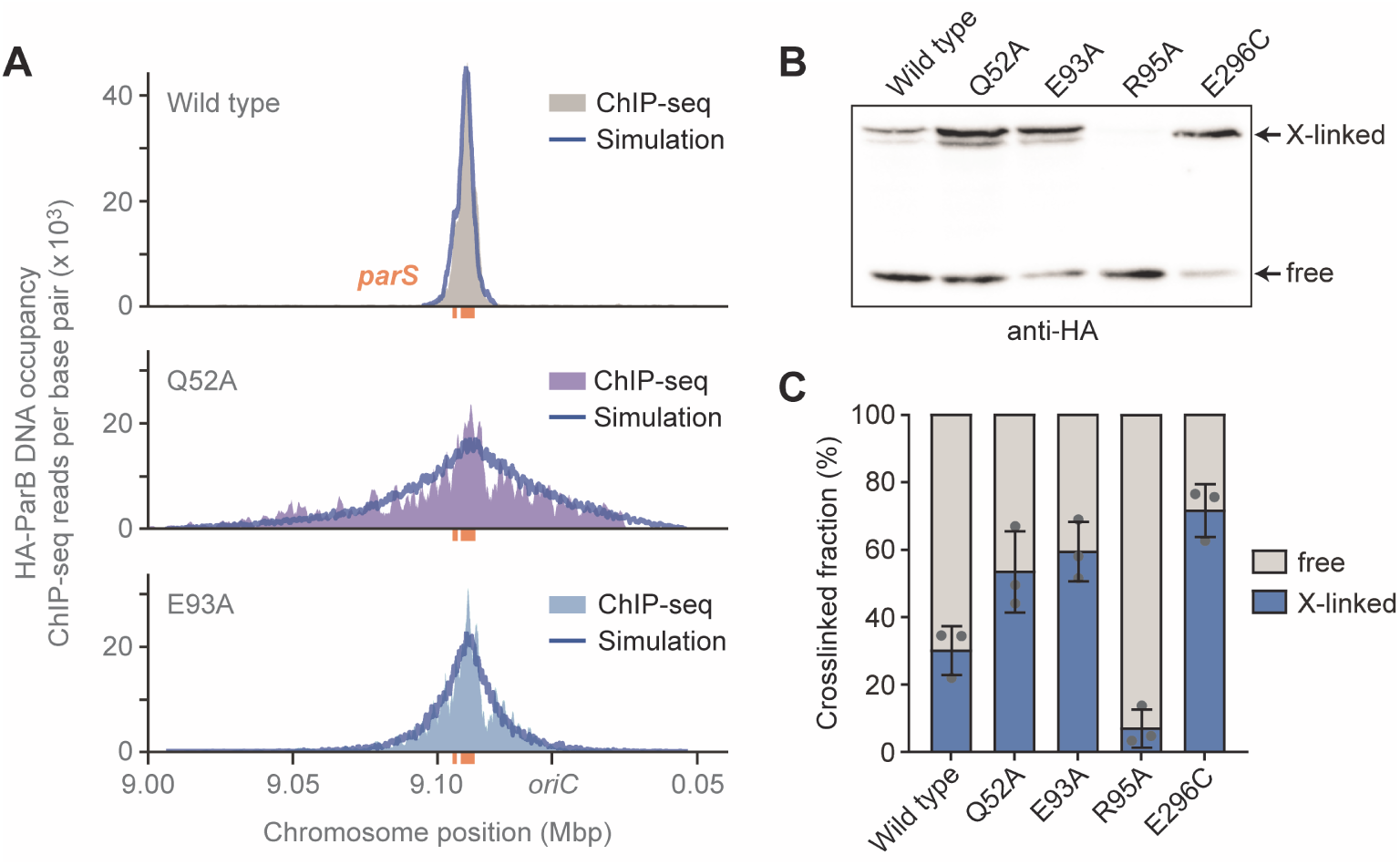
Catalytically inactive ParB variants spread over longer distances from *parS* sites *in vivo*. **(A)** ChIP-seq analysis of the spreading activity of different ParB variants. *M. xanthus* strains MO81 (wild type), MO82 (Q52A) and MO83 (E93A) were depleted of wild-type ParB (20 h) and concomitantly induced to produce the indicated HA-ParB variants. Cells were fixed with formaldehyde and subjected to ChIP-seq analysis with an anti-HA antibody. The graphs show the DNA occupancies plotted against the chromosomal position in an ∼200 kb region surrounding *oriC*. The results of stochastic 1D lattice simulations of the loading, sliding (diffusion) and dissociation of wild-type *Mx*ParB and its catalytically inactive variants are shown as blue traces that are superimposed on the ChIP-seq profiles. **(B)** *In vivo* site-specific crosslinking of selected HA-ParB variants. Cultures of strains MO85 (G37C), MO86 (G37C Q52A), MO87 (G37C E93A), JH5 (G37C R95A) and JHA5 (E296C) were grown as described in (A). After treatment with the thiol-specific crosslinker BMOE, cells were subjected to Western blot analysis with an anti-HA antibody. The signals produced by the crosslinked (X-linked) and non-crosslinked (free) species are indicated by arrows. **(C)** Quantification of the fraction of crosslinked and non-crosslinked species in the analysis shown in (B). Data represent the mean of three experiments (±SD). See also Figures S7A and S7B.

To further investigate the behavior of the different variants *in vivo*, we determined their opening state by site-specific *in vivo* crosslinking. For this purpose, *Mx*ParB variants additionally carrying an engineered cysteine residue (G37C) were synthesized in *M. xanthus* cells depleted of the wild-type protein. Subsequently, the cells were treated with BMOE, and crosslinking product were detected by Western blot analysis (**Figures 4B** and **4C**). In line with the *in vitro* results (**Figure 3A**), the Q52A and E93A variants showed a significantly higher fraction of crosslinked (closed) species than the wild-type protein. A variant unable to form rings (R95A), by contrast, was hardly detectable in the closed form. As a control, we also analyzed wild-type *Mx*ParB with a cysteine residue introduced at the C-terminal end of the constitutively associated dimerization domain (E296C). As observed *in vitro* (**Figure S6H**), it was crosslinked with high efficiency, further proving the validity of the crosslinking approach. Together, these results confirm that the extent of ParB spreading scales with the stability of the ring state *in vivo*.

As a test whether the sliding of ParB rings could indeed explain the distributions measured *in vivo*, we developed a stochastic 1D lattice simulation of the loading of ParB dimers at the 24 *parS* sites of *M. xanthus* and their subsequent sliding along the DNA. In the absence of any evidence to the contrary, we assumed that dimers undergo 1D Brownian diffusion along the DNA (as opposed to directed sliding) and that they cannot pass through one other. The degree of ParB spreading in such a model is described by a length-scale λ, representing the average distance that a ParB dimer would diffuse along the DNA, in the absence of obstacles, before dissociating. Comparing the results of the model against the ChIP-seq profiles (**Figure 4A**), we found good agreement for λ = 3.0 kb, which means that the experimental data is consistent with ParB dimers diffusing on average 3.0 kb before dissociating. As is clear from the profiles, the catalytically inactive Q52A and E93A variants spread much further, and we obtained the best agreement for λ = 43.8 kb and 14.1 kb, respectively. These results add to the idea that CTP hydrolysis may serve as a timing mechanism that limits the lifetime of sliding ParB rings, thereby confining their diffusion to the immediate vicinity of the *parS* cluster.

### Impairment of nucleotide hydrolysis reduces the mobility of *Mx*ParB within the cell

To further clarify the role of nucleotide hydrolysis in partition complex formation, we went on to study the dynamics of *Mx*ParB in different activity states using fluorescence-recovery-after-photobleaching (FRAP) analysis. For this purpose, wild-type *Mx*ParB and its Q52A and E93A variants were fused to the green fluorescent protein mNeonGreen (Shaner et al., 2013) and produced under the control of an inducible promoter in a conditional *parB* mutant. After depletion of the native *Mx*ParB protein, all fusion proteins formed distinct foci (**Figure 5A**), again confirming that CTP hydrolysis is not required for partition complex assembly (compare **Figure 4A**). As reported previously (Harms et al., 2013; Iniesta, 2014; Lin et al., 2017), the wild-type protein localized at a defined distance (∼1 µm) from the cell poles. When bleaching one of the two foci observed in pre-divisional cells, we found that the fluorescence signal recovered with a half-time of 20 ± 9 s, with the intensity of the second focus decreasing proportionally (**Figure 5A**, top). This result demonstrates that partition complexes are dynamic structures that exchange *Mx*ParB dimers in a rapid and continuous fashion, consistent with previous results on plasmid-encoded ParB proteins (Debaugny et al., 2018). Importantly, when the same analysis was performed with cells producing catalytically inactive *Mx*ParB variants, we observed no (Q52A) or only marginal (E93A) signal recovery (**Figure 5A**, middle and bottom). The excessive spreading of these proteins can thus be explained by a considerably longer residence time in the origin region, caused by a defect in ring opening.

**Figure 5.**
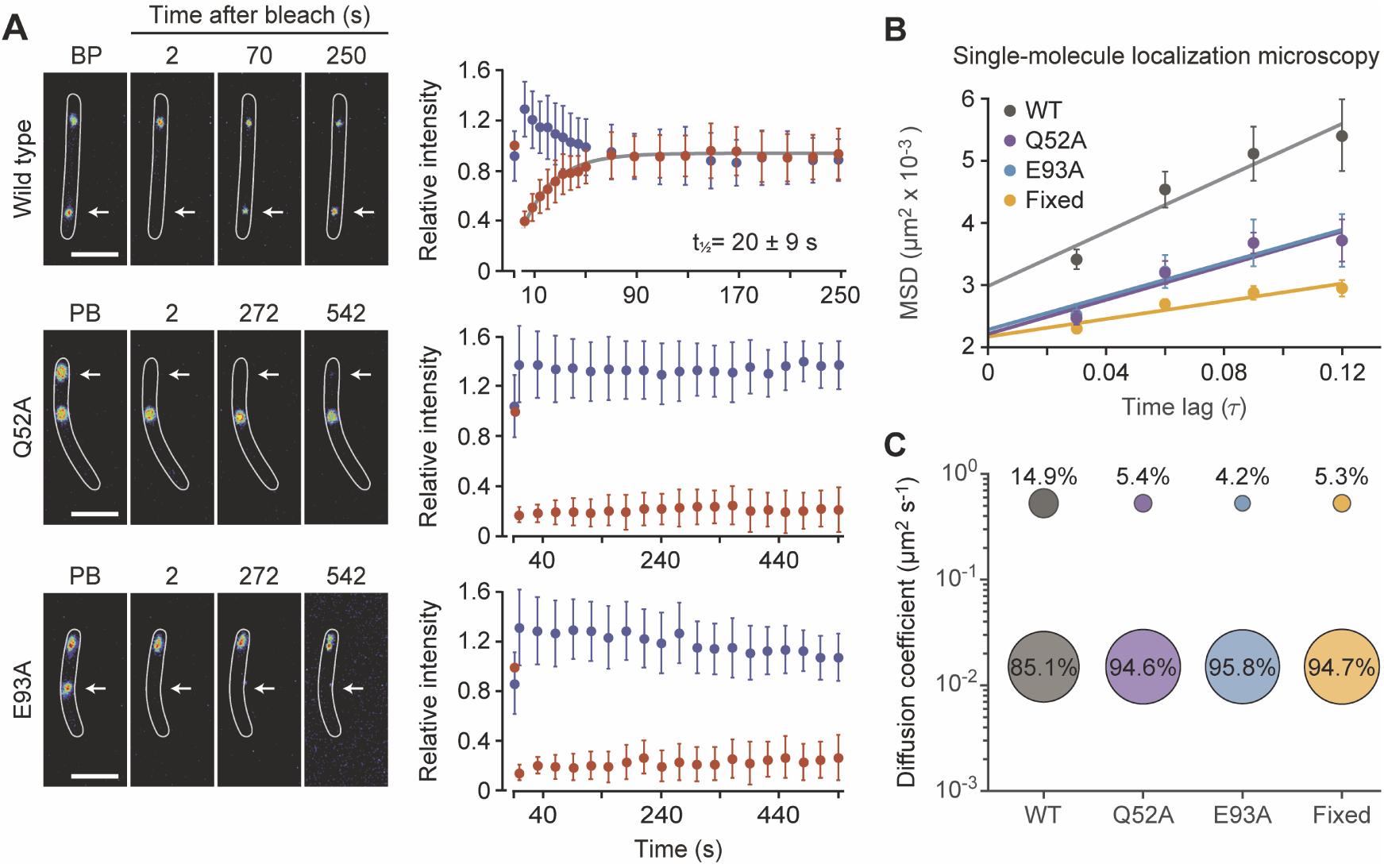
CTP hydrolysis determines the dynamics of partition complexes. **(A)** FRAP analysis of the mobility of mutant mNeonGreen-ParB variants. Strains MO78 (wild type), MO79 (E93A) and MO80 (Q52A) were depleted of wild-type *Mx*ParB (20 h) and concomitantly induced to the synthesize the indicated mNeonGreen-*Mx*ParB fusions. In cells containing two clearly distinguishable partition complexes, one of the foci was bleached by a short laser pulse, and the recovery of the signal was followed over time. The panels on the left show fluorescence images of representative cells before photobleaching (BP) and at the indicated times after application of the laser pulse. White arrows indicate the bleached region. Scale bars: 3 μm. The graphs show the average relative integrated intensities of the bleached (red) and unbleached (blue) partition complex (±SD) as a function of time (n= 38 cells for MO78, 16 cells for MO79 and 16 cells for MO80). The recovery half-time (± SD) for the wild-type protein was determined by fitting of the data to a single-exponential function (dark grey line). **(B)** Mean-squared displacement (MSD) analysis of the mobility of the indicated mNeonGreen-ParB variants, based on live-cell single-molecule localization microscopy data. Chemically fixed cells producing the wild-type fusion were analyzed as a reference. The graph shows the MSD (± SD) values for the first four time lags (τ) from all tracks containing more than four points as well as linear fits of the data points (colored line). **(C)** Bubble plot summarizing the diffusional behavior of the indicated mNeonGreen-ParB variants and a fixed reference sample. The values were obtained by fitting the probability distributions of the frame-to-frame displacements to a two-component Gaussian mixture model (GMM) assuming that the ParB dimer populations are composed of a mobile, open (top) and a static, closed (bottom) fraction. See also **Figure S7C**.

We hypothesized that the recovery rate measured for the wild-type protein is representative of the dissociation rate of *Mx*ParB rings from the DNA. Returning to the stochastic simulations described above, we fixed the dissociation half-time at the value suggested by FRAP analysis (20 s^-1^) and then used the model and the ChIP-seq profiles to predict the 1D diffusion coefficient of wild-type *Mx*ParB dimer diffusion along DNA. This analysis resulted in a value of (1.80 ± 0.06) x 10^-2^ μm^2^ s^-1^ (ranges here and below are 95% confidence intervals). Assuming that the catalytically inactive variants diffuse along DNA at the same rate as the wild-type protein, we can use this diffusion coefficient in simulations of the two variants to predict their dissociation half-times. The values obtained for the Q52A (4275 ± 4 s) and E93A (446 ± 2 s) proteins are 200 and 20 times higher than that for wild-type *Mx*ParB, respectively, consistent with the lack of recovery in the FRAP analysis (**Figure 5A**, middle and bottom). Indeed, such slow rates would be very challenging to measure experimentally, given cell growth and cell cycle progression.

Next, we performed single-particle tracking to clarify the mobility of *Mx*ParB in the DNA-loaded and released state. For this purpose, we again employed the above-described mNeonGreen fusions and followed their motion in cells depleted of the native *Mx*ParB protein. Calculating the average mean squared displacement of the entire population of dimers, we found that the wild-type protein was on average significantly more mobile than the Q52A and E93A variants (**Figure 5B**). A detailed analysis of the underlying single-particle tracks revealed that each variant showed two distinct diffusion regimes. The distribution of step sizes suggested the existence of both a mobile (*D* ∼ 0.54 ± 0.025 μm^2^ s^-1^) and a static (*D* ∼ 0.015 ± 0.001 μm^2^ s^-1^) dimer population, likely representing *Mx*ParB in the released (open) and the DNA-loaded ring state, respectively (**Figures 5C** and **S7C**). Notably, the diffusion coefficient of the static population was similar to what we predicted for DNA-associated *Mx*ParB rings based on our stochastic simulations above. For the wild-type protein, the mobile fraction comprised 15% of all dimers (**Figure 5C**), consistent with the rapid exchange of *Mx*ParB between partition complexes observed by FRAP analysis (compare **Figure 5A**). The proportion of mobile dimers decreased to ∼4-5% for the two catalytically inactive variants, reaching the background level measured for formaldehyde-treated, fixed cells. These results clearly demonstrate that ParB rings remain stably associated with the origin region in the absence of CTP hydrolysis, consistent with the larger fraction of closed rings observed in this condition (**Figures 4B** and **4C**).

### ParB CTPase activity is required for proper chromosome segregation *in vivo*

Despite their aberrant behavior, the Q52A and E93A variants formed well-defined, albeit more extensive, partition complexes (see **Figures 4** and **5A**), raising the question whether these assemblies were still able to support chromosome segregation. To address this issue, we produced wild-type or mutant *Mx*ParB-mNeonGreen fusions in cells depleted of native *Mx*ParB and then determined the localization patterns of the chromosomal origin regions, using the foci formed by these fusion proteins as a proxy (**Figure 6A**). Cells were imaged after a limited time of depletion (20 h) to ensure that potential functional defects of the tagged proteins did not yet affect their morphology, facilitating a comparison of the different strains. A demographic analysis showed that a strain producing the wild-type fusion protein exhibited a normal segregation pattern (Harms et al., 2013; Iniesta, 2014), with a single origin region in G1 phase and two well-separated and symmetrically arranged sister origin regions in late S- and G2-phase (**Figures 6A** and **6B**). In cells producing the Q52A and E93A variants, by contrast, the origin copies were poorly separated and mostly localized in the same half of the cell, indicating that the partition complexes formed were dysfunctional. A quantification of the number of fluorescent foci per cell (**Figure 6B**) and flow cytometry (**Figure 6C**) confirmed that this defect in origin segregation gave rise to daughter cells with abnormal chromosome content. Notably, upon prolonged incubation, cells producing the Q52A and E93A variants showed drastically reduced growth (**Figure 6D**), combined with cell lysis, and a severe decrease in fitness (**Figure 6E**). Thus, CTP hydrolysis is essential for ParB function.

**Figure 6.**
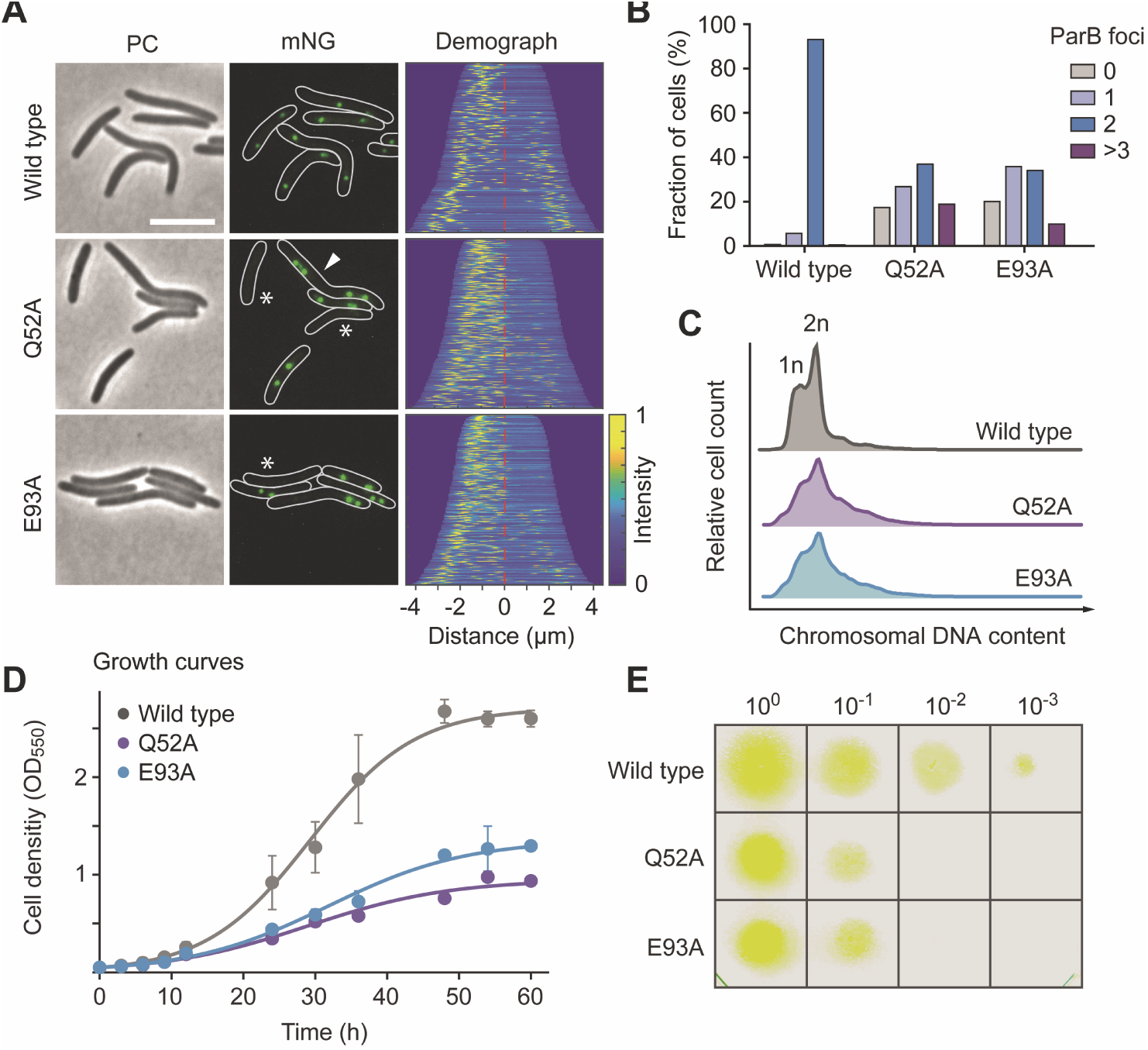
CTP hydrolysis is required for proper chromosome segregation. **(A)** Localization patterns of mNeonGreen-ParB variants. Strains MO78 (wild type), MO79 (E93A), and MO80 (Q52A) were depleted of wild-type ParB (20 h) and induced to produce the indicated mNeonGreen-ParB variants prior to analysis by fluorescence microscopy (scale bar: 5 µm). Asterisks indicate cells without partition complexes. The white arrowhead indicates a representative cell with multiple foci, indicative of a chromosome segregation defect. The demographs show the fluorescence profiles of a random subpopulation of cells sorted according to cell length and stacked on top of each other, with the shortest cell on top and the longest cell at the bottom (n = 219 cells for MO78, 291 cells for MO79 and 306 cells for MO80). **(B)** Quantification of the number of mNeonGreen-MxParB foci (partition complexes) in cells from the cultures analyzed in (A). **(C)** Chromosomal DNA content of MO78 (wild type), MO79 (E93A), and MO80 (Q52A) cells grown as described in (A). The cultures were treated with a DNA stain and analyzed by flow cytometry. Shown are histograms giving the distribution of fluorescence intensities in the different cell populations (n = 10,000 cells per strain). The fluorescence intensities corresponding to one (1n) or two (2n) chromosome equivalents are indicated. **(D)** Growth curves of strains MO78 (wild type), MO79 (E93A), and MO80 (Q52A). Cultures grown in the presence of CuSO_4_ to an OD_550_ of 0.8 were washed three times, diluted and cultivated in liquid medium supplemented with vanillate to produce the mutant ParB variants in place of the wild-type protein. Growth was then monitored by following the OD_550_ over a period of 60 h. The data shown represent the mean of two independent experiments (±SD). **(E)** Growth of the strains described in (D) on solid medium. Cells were grown in the presence of CuSO_4_ to an OD_550_ of 0.8, washed three times and serially diluted. 5 µl aliquots of the suspensions were then spotted onto CTT agar plates containing vanillate to induce the expression of the indicated mutant ParB variants in place of the wild type protein. The plates were incubated for 36 h before imaging.

## DISCUSSION

ParB acts as a DNA-sliding clamp whose loading and release is driven by a CTP-dependent conformational cycle. In this study, we shed light on thus-far poorly understood aspects of their biology, including the structural changes leading to ring closure, the mechanism of nucleotide hydrolysis and the role of their CTPase activity in partition complex dynamics and function. Our results close important gaps in the knowledge of this new class of molecular switches and provide a comprehensive basis for in-depth mechanistic studies of their conformational dynamics and their precise role in the DNA segregation process.

### The CTP-dependent conformational cycle of ParB

A key feature of ParB is its site-specific loading at *parS* sequences. It has been proposed that the interaction of the two DNA-binding domains with the palindromic *parS* motif juxtaposes the two NB domains and helps to overcome a kinetic barrier that otherwise prevents their interaction. In line with structural data on *B. subtilis* ParB and Noc (Jalal et al., 2021; Leonard et al., 2004; Soh et al., 2019), our HDX analyses suggest that this barrier may be a transition of the two ParB subunits from a compact apo to a ligand-bound loose state (**Figure 7**). The binding of either CTP or *parS* is sufficient to partially destabilize the compact state by inducing distinct conformational changes that interfere with the formation of the helix α4-α6 bundle. However, only in the presence of both ligands, the two ParB subunits adopt a conformation that enables productive face-to-face interactions between adjacent NB domains. Individual ParB subunits bind CTP with appreciable affinity, likely ensuring the saturation of their nucleotide-binding sites under physiological conditions. The need for additional structural changes induced by *parS* binding thus not only ensures the site-specific loading of ParB in the origin region but also prevents premature ring formation in the non-DNA-bound state. Notably, nucleotide hydrolysis is only observed in conditions that favor homodimerization of the NB domains. Thus, a functional catalytic site is only formed upon ring closure, even though the ParB/Srx module appears to be relatively static and all residues involved in nucleotide binding and catalysis are located in the *cis*-subunit. The critical role of the *trans*-subunit may be largely explained by its stabilizing effect on helix α4, which may facilitate interactions of residues E126, N127 and R130 with the triphosphate moiety, the Mg^2+^ ion and the water network that are critical for catalysis.

**Figure 7.**
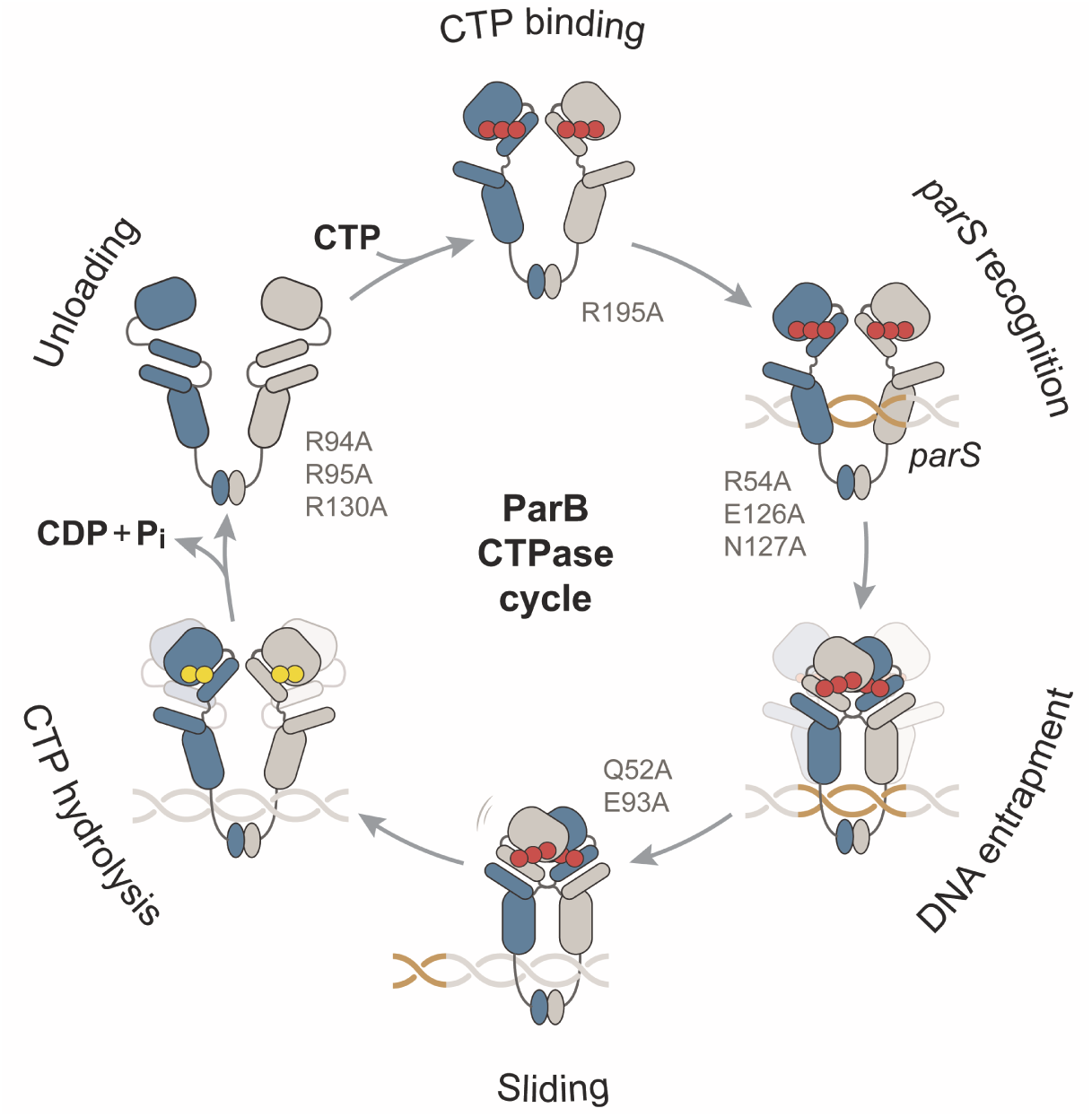
CTPase cycle of ParB. The interaction of apo ParB with CTP (red filled circles) destabilizes the packing of helices α4 and α5 and, thus, releases the CTP-bound NB domain. This structural change is reinforced by the association of the DNA-binding domain with a *parS* motif (highlighted in orange). DNA-binding allows a face-to-face interaction between the two adjacent NB domains, likely by reorienting helix α5 and thus facilitating its interaction with helix α4 of the *trans*-unit, respectively. Homodimerization of the NB-domains leads to ring closure and thus entraps the bound DNA in between the two arms of the ParB dimer. Moreover, it occludes and reorients the DNA-binding domain, thereby reducing the affinity of the closed ParB dimer for the *parS* sequence. As a consequence, the ParB ring slides away laterally from *parS*, which in turn enables the binding of another ParB dimer. The CTPase activity of ParB acts as a timing mechanism that controls the diffusion time of ParB rings on DNA. Nucleotide hydrolysis and the release of P_i_ destabilize the interaction between the two NB domains, leading to their dissociation and ring opening. CDP (yellow filled circles) has a low affinity for the NB domain and dissociates, thereby inducing a switch of the two subunits back to the apo form, followed by the unloading of the ParB dimer from the origin region. The open dimer then associates again with a *parS* site, thereby re-entering the cycle.

Our work shows that key driving forces behind nucleotide hydrolysis are linked to dynamic interactions between the active-site water network and the two catalytic residues Q52 and E93. Both of these residues are highly conserved in ParB homologs, whereas they are absent from the catalytically inactive NB domain of PadC. The precise catalytic mechanism still remains to be determined. However, it is conceivable that E93 serves as a general base, while Q52 may contribute to reducing the activation energy barrier by properly orienting waters in the network. In addition, Q52 may bind and stabilize the leaving phosphate to prevent the reverse reaction, *i.e.*, the formation of CTP. Interestingly, the active site of small GTPases from the Ras family also contains an essential glutamine residue, whose precise role in nucleotide hydrolysis has long been a matter of debate (Calixto et al., 2019; Kamerlin et al., 2013; Sprang, 1997). The absence of an obvious base in the immediate vicinity of the γ-phosphate led to the initial hypothesis that Q61 in p21^ras^ could act as the general base. However, later studies showed that GTP itself takes over this function in a substrate-assisted pathway (Schweins et al., 1995; Schweins et al., 1994), whereas Q61 may rather stabilize the transition state (Martin-Garcia et al., 2012). It is unlikely that Q52 of *Mx*ParB has a similar role, given its distance from the CTP γ-phosphate group. Our MD simulations show that the Q52 side chain shows considerably flexibility. Moreover, the active-site water network is dynamic and alternates between different compositional states. Together, these features may explain the low catalytic rate of *Mx*ParB.

In the post-hydrolysis complexes, the water network is displaced by the thiophosphate molecule generated during the catalytic process. Notably, this molecule interacts with the side chains of residues Q52 and R54 in the lid structure formed by loop α1/α2. It is tempting to speculate that, under physiological conditions, these interactions affect the dynamics of the lid and facilitate phosphate release. CTP hydrolysis and the loss of the phosphate group may change the configuration of helix α4, which could in turn destabilize its interaction with helices α5/α6 of the *trans*-subunit, thereby triggering the dissociation of the NB domains and ring opening. Consistent with this idea, we found that a mutation of residue N127, which interacts with the CTP γ-phosphate group as well as the released phosphate, completely abolishes MxParB ring formation. CDP has a low affinity for ParB (Osorio-Valeriano et al., 2019) and thus readily dissociates from the NB domains. As a consequence, ParB may transition to the apo conformation and leave the DNA (**Figure 7**). It then requires both CTP and *parS*-binding to return back to the DNA-bound closed state, ensuring its reloading close to *oriC*. Partition complexes thus represent dynamic self-organizing systems that use the free energy of CTP hydrolysis to establish a one-dimensional diffusion gradient of ParB rings on the chromosomal origin region, originating at *parS* and shaped by the CTPase activity of ParB. This behavior sets them apart from biomolecular condensates, which are self-assembling systems that form spontaneously without the need for continuous energy input (Alberti et al., 2019).

Interestingly, ParB rings can also be released from DNA in the absence of nucleotide hydrolysis, based on the spontaneous dissociation of the NB domains and subsequent loss of the nucleotides. The efficiency of this alternative pathway is inversely correlated with the concentration of free CTP during the opening event, i.e., the rate at which the lost nucleotides are replaced. The cellular concentration of CTP is in the range of 0.3 mM (Buckstein et al., 2008) and thus just high enough to saturate the nucleotide binding site of ParB. Under these conditions, nucleotide hydrolysis is the main driver of ring opening. However, it is conceivable that changes in the physiological state reduce the CTP level, thereby directly modulating the dynamics and function of partition complexes. This possibility to establish a direct regulatory link between cell physiology and chromosome segregation could explain the preference of ParB proteins for CTP, because the more commonly used cofactors ATP and GTP have considerably larger pool sizes and may thus provide less sensitivity to physiological changes.

### Role of CTPase activity in chromosome segregation

Our results show that the ParB CTPase activity is required for proper chromosome segregation *in vivo*, even though it is dispensable for the loading of ParB rings and the formation of spatially defined partition complexes. The reasons underlying this observation are still incompletely understood. In the absence of nucleotide hydrolysis, ParB remains stably associated with the origin region. Its longer diffusion times result in more extensive spreading, which increases the size of the partition complexes by up to fivefold (from 33 kb to 175 kb). Compared to the size of the chromosome (∼9.4 Mb), the fractional increase in DNA coverage is very small. However, it is possible that interactions between ParA and ParB need to be highly confined to enable a proper readout of the ParA gradient and ensure the unidirectional movement of the origin regions toward the cell poles. These interactions may be further compromised by the considerable decrease in the local concentration of ParB around the *parS* cluster that is observed under conditions of increased spreading. Furthermore, the dynamic conformational changes driven by CTP binding and hydrolysis may not only control the size of partition complexes but also have a more direct role in the segregation process. The docking site for ParA is thought to reside in the N-terminal non-structured region of ParB (Ah-Seng et al., 2009; Figge et al., 2003; Leonard et al., 2005; Radnedge et al., 1998; Surtees and Funnell, 1999). Recent work has suggested that the ability of these regions to interact with ParA is modulated by the opening state of ParB dimers (Taylor et al., 2021), suggesting that the lifetime of ParB rings could have an immediate effect on the dynamics of origin movement. The kinetics of ring opening may also affect the DNA-condensing activity observed for some ParB homologs, which is thought to involve bridging interactions between distal ParB dimers (Broedersz et al., 2014; Debaugny et al., 2018; Graham et al., 2014; Sanchez et al., 2013; Song et al., 2017) or non-specific DNA-binding by the C-terminal dimerization domain (Fisher et al., 2017; Taylor et al., 2015). Finally, partition complexes mediate, directly or indirectly, the loading of the SMC/condensin complex (Böhm et al., 2020; Gruber and Errington, 2009; Sullivan et al., 2009; Tran et al., 2017), which also contributes to chromosome segregation in *M. xanthus* (Anand et al., 2020). Previous work in *Corynebacterium glumaticum* has shown that a ParB variant with an amino acid exchange equivalent to R94A in *M. xanthus* (compare **Figure S6C**) still recruits SMC to the *parS* sites but no longer supports its translocation across the chromosome (Böhm et al., 2020). This finding suggests that the CTP-dependent formation of ParB rings at *parS* and their spreading on DNA is critical for proper SMC recruitment. However, the exact role of the ParB CTPase cycle in this process still remains to be determined.

## Conclusions

ParB proteins are widely conserved among prokaryotes and represent an ancient family of nucleotide-dependent molecular switches with critical functions in DNA segregation and other fundamental cellular processes. The unique characteristic of this prototypic novel family of regulatory NTPases is their preference of CTP over the purine nucleotides ATP and GTP, mediated by their N-terminal ParB/Srx module. Our work now clarifies the CTPase cycle of ParB proteins and, thus, provides detailed insight into the mechanistic basis and physiological functions of their switch-like behavior. Our characterization of the catalytic activity of ParB sheds new light onto the mechanisms of nucleotide hydrolysis and nucleotide-dependent regulation and offers new perspectives in the investigation of nucleotide-dependent regulation in biology. Importantly, there are many non-ParB proteins with domains homologous to the NB domain, and it will be interesting to analyze their functions and determine whether they use a CTP-dependent switch mechanism similar to that identified for ParB.

## ACKNOWLEDGEMENTS

We thank Julia Rosum for excellent technical assistance, Paul Weiland for help with the structural analysis as well as Adam Jalal and Tung Le for sharing unpublished results. Moreover, we acknowledge the European Synchrotron Radiation Facility (ESRF, Grenoble, France) for assistance in data collection collection and Silvia G. Sierra from the Flow Cytometry and Imaging Facility of the Max Planck Institute for Terrestrial Microbiology (Marburg) for help with the flow cytometry analysis. This work was funded by the Max Planck Society (Max Planck Fellowship to M.T.; Core funding to S.M.), the German Research Foundation (DFG) (Project 269423233 – TRR 174 to M.B., G.B. and M.T.; German Excellence Strategy – EXC 2033 – 390677874 – RESOLV to L.V.S.; grant BR 2915/6-2 to M.B.; DFG Core Facility for Interactions, Dynamics and Macromolecular Assembly, project 324652314, to G.B.), the Swiss National Science Foundation (grant 31003A_182576 to P.H.V.) and the European Union’s Horizon 2020 research and innovation program (Marie Sklodowska-Curie grant, agreement No 801459 to C.K.D; Marie Sklodowska-Curie grant, agreement No 659174 to L.C-G.).

## AUTHOR CONTRIBUTIONS

M.O.V. and M.T. conceived the study. M.O.V. and J.H. conducted the biochemical and cell biological studies. F.A. performed the crystallization screens and solved the crystal structures reported in this paper. C.K.D. conducted the molecular dynamics simulations. W.S. performed the HDX experiments. G.P. performed the ChIP-seq experiments. L.C. and S.M. conducted the stochastic simulations. G.G. and H.F. performed the single-particle tracking studies. L.C.-G. performed the FRAP experiments. P.G. assisted with the ITC experiments. M.O.V., F.A., C.K.D., W.S., G.P., L.C., G.G., H.F., L.C.-G., M.B., P.V., S.M., L.S., G.B., and M.T. analyzed the data. M.B., P.V., S.M., L.S., G.B. and M.T. secured funding and supervised the study. M.O.V. and M.T. wrote the paper, with input from all other authors.

## METHODS

### LEAD CONTACT AND MATERIALS AVAILABILITY

Further information and requests for resources and reagents should be directed to and will be fulfilled by the Lead Contact, Martin Thanbichler. All plasmids and strains generated in this study are available from the Lead Contact without restriction.

### EXPERIMENTAL MODEL AND SUBJECT DETAILS

#### Media and growth conditions

*M. xanthus* DK1622 and derivative strains were grown at 32 °C in CTT medium (Hodgkin and Kaiser, 1977), supplemented with kanamycin (50 µg/mL) or oxytetracycline (10 µg/mL) when appropriate. *E. coli* strains were cultivated at 37 °C in LB medium containing antibiotics at the following concentrations (µg/mL in liquid/solid medium): ampicillin (100/200), chloramphenicol (20/34), kanamycin (30/50). The expression of genes placed under the control of the P*_van_* (Iniesta et al., 2012) or P*_cuoA_* (Gomez-Santos et al., 2012) promoter was achieved by supplementation of the media with 0.5 mM sodium vanillate or 0.3 mM CuSO_4_, respectively. Growth curves were fitted to a previously described model (Huang, 2011) using the Solver function of Microsoft Excel 2019.

### METHOD DETAILS

#### Plasmid and strain construction

The construction of bacterial strains and plasmids is detailed in **Tables S3** and **S4**. Oligonucleotides used for cloning purposes are listed in **Table S5**. *E. coli* TOP10 (Invitrogen) was used as host for cloning purposes. All plasmids were verified by DNA sequencing. *M. xanthus* was transformed by electroporation (Kashefi and Hartzell, 1995). Non-replicating plasmids were integrated into the chromosome by site-specific recombination at the phage Mx8 *attB* site (Magrini et al., 1999) or single-homologous recombination at the MXAN_0018/0019 locus (Iniesta et al., 2012). Gene replacement was achieved by double-homologous recombination using the counter-selectable *galK* marker (Ueki et al., 1996). Proper chromosomal integration or gene replacement was verified by colony PCR.

#### Widefield fluorescence microscopy

*M*. *xanthus* strains carrying the endogenous *parB* gene under the control of a Cu^2+^-inducible promoter and the indicated *mNeonGreen*-*parB* variants under the control of a vanillate-inducible promoter were pre-grown in CTT medium containing CuSO_4_, washed three times with fresh CTT medium, and then cultivated for 20 h in CTT medium supplemented with vanillate to deplete the wild-type protein and concomitantly produce the fluorescently tagged ParB variants.

To record still images, cells were spotted on 1.5% agarose pads in TPM buffer (10 mM Tris/HCl, 8 mM MgSO_4_, 1 mM potassium phosphate, pH 7.6) supplemented with 10% CTT medium (corresponding to a final content of 0.2% casitone) (Schumacher and Søgaard-Andersen, 2018). Images were taken with a Zeiss Axio Imager.M1 microscope equipped with a Zeiss Plan Apochromat ×100/1.40 Oil DIC objective and a pco.edge 3.1 sCMOS camera (PCO) or with a Zeiss Axio Imager.Z1 microscope equipped with a ×100/1.46 Oil DIC objective and a pco.edge 4.2 sCMOS camera (PCO). An X-Cite 120PC metal halide light source (EXFO, Canada) and ET-YFP or ET-TexasRed filter cubes (Chroma, USA) were used for fluorescence detection. Images were recorded with VisiView 4.0.0.14 (Visitron Systems) and processed with Metamorph 7.7.5 (Molecular Devices) and Adobe Illustrator CS6 (Adobe Systems). The subcellular distribution of fluorescence signals was analyzed with BacStalk.

FRAP analysis was performed with the Zeiss Axio.Observer Z1 setup described above, using a 488 nm-solid state laser and a 2D-VisiFRAP multi-point FRAP module (Visitron Systems, Germany), with 396-ms pulses at a laser power of 15%. After acquisition of a pre-bleach image and application of a laser pulse, 10 images were taken at 6 s intervals, followed by 10 images at 20 s intervals (strain MO78). Alternatively, two images were taken at 6 s intervals, followed by 18 images at 30 s intervals (strains MO79 and MO80). For each time point, the integrated fluorescence intensities of the whole cell, the bleached region and an equally sized unbleached region were measured using Fiji 1.49 (Schindelin et al., 2012). Recovery half-times were calculated independently for every cell as described previously (Corrales-Guerrero et al., 2020) by fitting the data to a single-exponential function in MATLAB R2014b (Mathworks, USA).

#### Single-molecule localization microscopy (SMLM)

SMLM analysis was performed with an Elyra 7 (Zeiss) inverted microscope equipped with two pco.edge sCMOS 4.2 CL HS cameras (PCO AG) connected through a DuoLink (Zeiss), only one of which was used in this study. Cells were grown as described for widefield fluorescence microscopy and observed through an alpha Plan-Apochromat 63x/1.46 Oil Korr M27 Var2 objective in combination with an Optovar 1.6x (Zeiss) magnification changer, yielding a pixel size of 63 nm. During image acquisition, the focus was maintained with the help of a Definite Focus.2 system (Zeiss). For each time-lapse series, 3,300 frames were taken with 30 ms exposure time. Fluorescence was excited with 405 nm (50 mW) and 488 nm (100 mW) diode lasers, and signals were observed through a multiple beam splitter (405/488/561/641 nm) and laser block filters (405/488/561/641 nm) followed by a Duolink SR QUAD (Zeiss) filter module (secondary beam splitter: LP640, emission filter: BP 495-590). Cells were illuminated with the 488 nm laser (40% intensity), supported by the 405 nm laser (0.1% intensity) to increase the chance of dark-state mNeonGreen reverting back to a fluorescent state without affecting its fluorescence.

For single-particle tracking, spots were identified with the LoG Detector of TrackMate v6.0.1 (Tinevez et al., 2017), implemented in Fiji 1.53h (Schindelin et al., 2012), with an estimated diameter of 0.315 µm, a threshold of 6, and median filter and sub-pixel localization activated. To limit the detection of particles to single-molecules and avoid the necessity to pre-bleach mNeonGreen molecules, spots with a maximum intensity ≥ 200 were discarded. This threshold value was determined through analysis of the intensity distribution of all recorded mNeonGreen particles in a typical SMLM experiment. Spots were merged into tracks via the Simple LAP Tracker of TrackMate, with a maximum linking distance of 300 nm, no frame gaps allowed, and a minimum track length of 5, yielding a minimum n of total tracks per sample of 1800. To identify differences in protein mobility and in the proportions of mobile and static molecules between strains, the resulting tracks were subjected to mean-squared-displacement (MSD) and Gaussian-mixture-model (GMM) analysis in SMTracker 2.0 (Rosch et al., 2018) as described previously (Hernandez-Tamayo et al., 2021; Rosch et al., 2018). The average MSD was calculated for five separate time points per strain (τ = 30, 60, 90, 120 and 150 ms), followed by fitting of the data to a linear equation. The last time point of each track was excluded to avoid track-ending-related artifacts. GMM analysis was based on a double-Gaussian fit assuming two protein populations.

#### Protein purification

For the purification of His_6_-SUMO-ParB, *E. coli* Rosetta(DE3)pLysS cells carrying plasmid pMO104 were grown at 37 °C in 3 L of LB medium supplemented with ampicillin (200 µg/mL) and chloramphenicol (34 µg/mL). At an OD_600_ of 0.6, the culture was chilled to 18 °C and protein synthesis was induced by the addition of 1 mM IPTG prior to incubation of the cells overnight at 18°C. The cultures were harvested by centrifugation at 10,000 ×g for 20 min at 4 °C, washed with buffer ParB1 (25 mM HEPES/NaOH, pH 8.0, 300 mM NaCl, 0.1 mM EDTA, 5 mM MgCl_2_), and resuspended in buffer ParB2 (25 mM HEPES/NaOH, pH 8.0, 300 mM NaCl, 0.1 mM EDTA, 5 mM MgCl_2_) supplemented with 20 mM imidazole, 10 μg/mL DNase I and 100 μg/mL PMSF. After three passages through a French press (16,000 psi), the cell lysate was clarified by centrifugation (30,000 × g, 30 min, 4 °C), and the supernatant was subjected to immobilized-metal affinity chromatography (IMAC) using a 5 mL HisTrap HP column (GE Healthcare) equilibrated with buffer ParB1 containing 20 mM imidazole. Protein was eluted with a linear gradient of 20 to 250 mM imidazole in buffer ParB1 at a flow rate of 1 mL/min. Fractions containing high concentrations of ParB were pooled and dialyzed against 3 L of buffer ParB3 (25 mM HEPES, pH 7.6, 150 mM NaCl, 0.1 mM EDTA, 5 mM MgCl_2_, 10% (v/v) glycerol). After the addition of Ulp1 protease (Marblestone et al., 2006) and dithiothreitol (1 mM), the protein was incubated for 4 h at 4°C to cleave off the His_6_-SUMO tag. The solution was centrifuged for 30 min at 38,000 × g and 4°C to remove precipitates and applied onto a 5 mL HisTrap HP column (GE Healthcare) equilibrated with buffer ParB3. ParB was recovered from the flow-through, whereas His_6_-SUMO was retained on the column. The fractions containing ParB were concentrated in a spin concentrator (Amicon, MWCO 10,000). After the removal of precipitates by centrifugation at 30,000 ×g for 30 min, ParB was further purified by size exclusion chromatography (SEC) on a HighLoad 16/60 Superdex 200 pg column (GE Healthcare) equilibrated with buffer ParB3. Fractions containing pure protein were pooled and concentrated. After the removal of precipitates by centrifugation at 30,000 x g, the protein solution was snap-frozen in liquid N_2_ and stored at −80 °C until further use. Mutant ParB variants were purified essentially as described for the wild-type protein.

#### Crystallization and structure determination

Proteins were crystallized with the sitting-drop method at 20 °C in 0.5 µL drops consisting of equal parts of protein and precipitation solution. Wild-type ParB·CDP·P_S_ crystallized at a concentration of 150 mg/ml in a buffer containing 10% (w/v) glycerol, 0.2 M MgCl_2_, 0.1 M Tris/HCl pH 8.5 and 25 % (v/v) 1,2-propanediol, supplemented with 10 mM CTPγS after 48 h of incubation. ParB_Q52A_·CTPγS crystallized at a concentration of 150 mg/ml in a buffer containing 0.1 M Tris/HCl pH 8.5 and 20% (w/v) PEG 1000, supplemented with 10 mM CTPγS after 24 h of incubation. ParB_E93A_·CDP·P_S_ crystallized at a concentration of 150 mg/ml in a buffer containing 0.1 M phosphate-citrate buffer pH 4.2 and 40% (v/v) PEG 300, supplemented with 10 mM CTPγS after 24 h of incubation. Prior to data collection, crystals were cryo-protected with the respective mother liquor supplemented with 25% (v/v) glycerol. Datasets were collected under cryogenic conditions at the BESSY II beamline BL14.1 (Gerlach et al., 2016) and at EMBL beamline P14 at the PETRA III storage ring {McCarthy, 2018 #140), respectively. Phase-determination of ParB was achieved by molecular replacement using the structure of the *B. subtilis* ParB·CDP complex (PDB: 6SDK) (Soh et al., 2019) as a model. The phasing of ParB_Q52A_·CTPγS and ParB_E93A_·CDP·P_S_ was performed by molecular replacement using the wild-type ParB structure as a search model. The data were processed with XDS (Kabsch, 2010) and scaled with XSCALE (Kabsch, 2010). The substructure was determined with PHENIX-implemented Phaser and refined with PHENIX-refine prior to manual model building with Coot (Adams et al., 2010; Emsley and Cowtan, 2004). Final model validation was performed with the PDB-Redo Server (Joosten et al., 2014).

Protein structures were visualized with PyMOL 2.1 (www.pymol.org). LIGPLOT (Wallace et al., 1995) was used to generate protein-ligand interaction 2D maps. Protein domains were predicted using the PFAM database (El-Gebali et al., 2019).

#### Biolayer interferometry

Biolayer interferometric analyses were performed with a BLItz system equipped with High Precision Streptavidin (SAX) Biosensors (ForteBio). A double-biotinylated DNA fragment (234 bp) containing a single *parS* site was PCR-amplified from *M. xanthus* chromosomal DNA with primers carrying a 5′-biotin-triethylene glycol (TEG) group (BioTEG-*parS*-3-for/BioTEG-*parS*-3-rev). After its immobilization on the sensor surface and the establishment of a stable baseline, the association of ParB or its mutant variants (10 µM) was monitored in the presence of the indicated nucleotides. At the end the association step, the sensor was transferred into protein-free buffer containing or lacking nucleotides to follow the dissociation kinetics. The extent of non-specific binding was assessed by monitoring the interaction of the analytes with unmodified sensors. All analyses were performed in BLItz binding buffer (25 mM HEPES/KOH pH 7.6, 100 mM KCl, 10 mM MgSO_4_, 1 mM DTT, 10 µM BSA, 0.01% Tween 20), supplemented with nucleotides when indicated.

#### Isothermal titration calorimetry (ITC)

Nucleotide binding to ParB was measured using a MicroCal PEAQ-ITC system (Malvern Panalytical). ParB and its mutant derivatives were dialyzed extensively against ITC buffer (25 mM HEPES/NaOH, pH 7.6, 150 mM NaCl, 0.1 mM EDTA, 5 mM MgCl_2_). Nucleotides (CDP and CTPγS) were dissolved in the same buffer. Proteins (150 µM) were titrated with 13 consecutive injections (2 µL) of CTP or CTPγS stock solutions (1.25 mM) at 25 °C and 150 s intervals, with a duration of each injection of 4 s. The mean enthalpies of dilution were subtracted from the raw titration data before analysis. Titration curves were fitted to a one-set-of-sites model using the MicroCal PAEQ-ITC analysis software (Malvern Panalytical).

#### Nucleotide hydrolysis assays

Nucleotide hydrolysis was measured using a coupled enzyme assay (Ingerman and Nunnari, 2005; Kiianitsa et al., 2003). Reactions contained 4 µM wild-type or mutant ParB, 20 U/mL pyruvate kinase (Sigma Aldrich), 20 U/mL L-lactate dehydrogenase (Sigma Aldrich), 800 µg/mL NADH and 3 mM PEP in 200 µL reaction buffer (25 mM HEPES/KOH pH 7.4, 100 mM KCl, 10 mM MgSO_4_, 1 mM DTT). A DNA stem-loop (54 bases; *parS*-Mxan-wt) containing a wild-type *parS* site (250 nM) was added when indicated. After incubation for 10 min at 30°C, 150 µL of the reaction mixtures were transferred into a 96-well microtiter plate and supplemented with CTP to a final concentration of 1 mM to start the reactions. CTP hydrolysis was followed by measuring the decrease in NADH absorbance at 340 nm at 2 min intervals. Initial velocities were calculated by linear regression analysis of each time course and corrected for spontaneous CTP hydrolysis and NADH oxidation.

#### *In vitro* crosslinking

Mutant ParB variants carrying an additional G37C substitution were incubated with 5 mM tris(2-carboxyethyl)phosphine (TCEP) for 30 min at room temperature. TCEP was removed with a PD SpinTrap G-25 desalting column (GE Healthcare) equilibrated with reaction buffer (25 mM HEPES pH 7.6, 150 mM NaCl, 5 mM MgCl_2_, 0.1 mM EDTA). Proteins (10 µM) were mixed with the indicated concentrations of CTP or CTPγS and/or a *parS*-containing DNA hairpin (1 µM; *parS*-Mxan-wt) and preincubated for 5 min at room temperature. Subsequently, aliquots were taken from the reactions at the indicated time points and incubated with 1 mM bismaleimidoethane (BMOE) for 5 min at room temperature. The crosslinking reactions were quenched by the addition of dithiothreitol-containing SDS sample buffer. Samples were loaded onto an SDS-PAGE gel and protein was stained with Coomassie Brilliant Blue R-250. The intensities of the different bands were quantified with a ChemiDoc MP imaging system (Bio-Rad).

#### *In vivo* crosslinking

*M*. *xanthus* strains carrying the endogenous *parB* gene under the control of a Cu^2+^-inducible promoter and the indicated *HA*-*parB* variant harboring an additional G37C mutation under the control of a vanillate-inducible promoter were grown in CTT medium containing CuSO_4_, washed three times with frest CTT medium, and then cultivated for 20 h in CTT medium supplemented with vanillate to produce the mutant ParB variants in place of the wild-type protein. Cells were harvested, washed with cold TMP buffer (10 mM Tris/HCl pH 7.6, 1 mM sodium phosphate pH 7.6 and 8 mM MgSO_4_) and resuspended in 250 µL of cold TMP buffer with 0.1% glycerol (TMPG) to a final OD_550_ of 10. Crossliking was achieved by incubation of the cells with 2.5 mM BMOE for 5 min at room temperature. After quenching of the reaction with 10 mM DTT, the cells were collected by centrifugation and resuspended in SDS sample buffer. The cell lysates were subjected to SDS-PAGE and analyzed by Western blotting using an anti-HA primary antibody (rabbit; Millipore, #15-902R), an HRP-conjugated goat anti-rabbit secondary antibody (PerkinElmer) and the Western Lightning ECL Pro kit (PerkinElmer). The signals were quantified with a ChemiDoc MP imaging system (Bio-Rad).

#### Hydrogen-deuterium exchange (HDX) mass spectrometry

HDX-MS experiments on *Mx*ParB were carried out as described previously (Osorio-Valeriano et al., 2019) with minor modifications. The samples contained 27 µM *Mx*ParB in a buffer composed of 25 mM HEPES/KOH pH 7.6, 100 mM KCl and 10 mM MgSO_4_. Nucleotides (CDP, CTP, CTPγS) and *parS* DNA (*parS*-Mxan-wt) were used at final concentrations of 10 mM and 27 µM, respectively. The preparation of the HDX reactions was aided by a two-arm robotic autosampler (LEAP technologies). 7.5 μL of *Mx*ParB solution (with or without nucleotides and/or DNA) were mixed with 67.5 μL of D_2_O-containing buffer (25 mM HEPES/KOH pH 7.6, 100 mM KCl, 10 mM MgSO_4_) to start the exchange reaction. After 10, 30, 95, 1,000 and 10,000 s of incubation at 25 °C, 55 μL samples were taken from the reaction and mixed with an equal volume of quench buffer (400 mM KH_2_PO_4_/H_3_PO_4_, 2 M guanidine-HCl, pH 2.2) kept at 1 °C. 95 µL of the resulting mixture were injected into an ACQUITY UPLC M-Class System with HDX Technology (Waters) (Wales et al., 2008) through a 50 µL sample loop. Undeuterated samples of *Mx*ParB were prepared similarly by 10-fold dilution of the protein solution into H_2_O-containing buffer. The samples were flushed out of the loop with water + 0.1 % (v/v) formic acid (100 µL/min) over 3 min and injected to a column (2 mm x 2 cm) that was packed with immobilized protease type XIII from *Aspergillus saitoi* (Cravello et al., 2003) and kept at 12 °C for proteolytic digestion. The resulting peptides were collected on a trap column (2 mm x 2 cm) filled with POROS 20 R2 material (Thermo Scientific) kept at 0.5 °C. Subsequently, the trap column was placed in line with an ACQUITY UPLC BEH C18 1.7 μm 1.0 x 100 mm column (Waters), and the peptides were eluted at 0.5 °C using a gradient of water + 0.1 % (v/v) formic acid (A) and acetonitrile + 0.1 % (v/v) formic acid (B) at a flow rate of 30 μL/min as follows: 0-7 min/95-65 % A, 7-8 min/65-15 % A, 8-10 min/15 % A, 10-11 min/5 % A, 11-16 min/95 % A. After ionization of the peptides with an electrospray ionization source (250 °C capillary temperature, 3.0 kV spray voltage), mass spectra were acquired in positive ion mode over a range of 50 to 2000 *m/z* on a G2-Si HDMS mass spectrometer with ion mobility separation (Waters), using Enhanced High Definition MS (HDMS^E^) or High Definition MS (HDMS) mode for undeuterated and deuterated samples, respectively (Geromanos et al., 2009; Li et al., 2009). [Glu1]-Fibrinopeptide B standard (Waters) was employed for lock mass correction. During each chromatographic run, the protease type XIII column was washed three times with 80 µL of 4 % (v/v) acetonitrile and 0.5 M guanidine-HCl, and blank injections were performed between each sample. All measurements were carried out in triplicate.

*Mx*ParB peptides were identified from the non-deuterated samples (aquired with HDMS^E^) with the software ProteinLynx Global SERVER (PLGS, Waters), using low energy, elevated energy and intensity thresholds of 300, 100 and 1,000 counts, respectively. Peptides were matched using a database containing the amino acid sequence of *Mx*ParB and its reverse sequence. The search parameters were chosen as described previously (Osorio-Valeriano et al., 2019). After automated data processing by the software DynamX (Waters), all spectra were inspected manually and, if necessary, peptides were omitted (e.g. in case of a low signal-to-noise ratio or the presence of overlapping peptides).

#### Chromatin immunoprecipitation coupled to deep sequencing (ChIP-seq) analysis

*M. xanthus* strains carrying the endogenous *parB* gene under the control of a Cu^2+^-inducible promoter and *HA-parB* (MO81), *HA-parB_Q52A_* (MO82), *HA-parB_E93A_* (MO83) or *HA-parB_R95A_* (MO84) under the control of a vanillate-inducible promoter were grown overnight in the presence of CuSO_4_. The cells were washed three times in CTT medium and cultivated for an additional 24 h in 80 ml CTT medium supplemented with vanillate to deplete wild-type ParB and induce the synthesis of the HA-tagged ParB variants. At an OD_550_ of 0.5, the cultures were supplemented with 10 mM sodium phosphate buffer (pH 7.6) and then treated with formaldehyde (1% final concentration) at room temperature for 10 min to achieve crosslinking. After incubation for an additional 30 min on ice and three washes in phosphate-buffered saline (PBS, pH 7.4), the cells were harvested by centrifugation and stored at −80°C until further use. After resuspension in TES buffer (10 mM Tris/HCl pH 7.5, 100 mM NaCl, 1 mM EDTA), the cells were incubated for 10 min at 37°C in the presence of Ready-Lyse lysozyme solution (Epicentre) according to the manufacturer’s instructions. The lysates were then sonicated (Bioruptor® Pico, Diagenode) at 4°C using 15 bursts of 30 sec to shear DNA fragments to an average length of 0.2-0.5 kbp and cleared by centrifugation for 2 min at 14,000 rpm and 4°C. The volume of the lysates was adjusted (relative to the protein concentration) to 1 ml using ChIP buffer (16.7 mM Tris/HCl pH 8.1, 167 mM NaCl, 0.01% SDS, 1.1% Triton X-100, 1.2 mM EDTA) containing protease inhibitors (Roche) and pre-cleared with 80 μL of Protein-A agarose (Roche) and 100 μg BSA. 2% of each pre-cleared lysate were reserved as total input (negative control) samples. The pre-cleared lysates were then incubated overnight at 4°C with monoclonal rabbit Anti-HA Tag antibodies (Millipore, clone 114-2C-7) at 1:400 dilutions. Immunocomplexes were captured by incubation with Protein-A agarose beads (pre-saturated with BSA) for 4 h at 4°C. After collection by centrifugation, the beads were washed with low-salt washing buffer (20 mM Tris/HCl pH 8.1, 150 mM NaCl, 2 mM EDTA, 0.1% SDS, 1% Triton X-100), with high-salt washing buffer (20 mM Tris/HCl pH 8.1, 500 mM NaCl, 2 mM EDTA, 0.1% SDS, 1% Triton X-100), with LiCl washing buffer (10 mM Tris/HCl pH 8.1, 0.25 M LiCl, 1 mM EDTA, 1% NP-40, 1% deoxycholate,) and finally twice with TE buffer (10 mM Tris/HCl pH 8.1, 1 mM EDTA). The immuno-complexes were eluted from the Protein-A agarose beads with two times 250 μL elution buffer (0.1 M NaHCO_3_, 1% SDS; freshly prepared) and, in parallel to the total input samples, incubated overnight with 300 mM NaCl at 65°C to reverse the crosslinks. The samples were then treated with 2 μg of proteinase K for 2 h at 45°C in 40 mM EDTA and 40 mM Tris/HCl (pH 6.5). DNA was extracted using phenol:chloroform:isoamyl alcohol (25:24:1), ethanol-precipitated using 20 μg of glycogen as a carrier and resuspended in 50 μL of DNAse/RNAse-free water.

Immunoprecipitated chromatin (two biological replicates per condition) was used to prepare sample libraries for deep-sequencing (Fasteris SA, Geneva, Switzerland). ChIP-seq libraries were prepared using the DNA Sample Prep Kit (Illumina) following the manufacturer’s instructions. Single-end runs with 50 cycles were performed on an NextSeq High (Illumina) next-generation DNA sequencing instrument, yielding several million reads per sequenced samples. The single-end sequence reads, stored in FastQ files, were mapped against the genome of *M. xanthus* DK1622 (GenBank: CP000113.1) using Bowtie 2 v2.3.4 + galaxy0, available through the web-based analysis platform Galaxy (https://usegalaxy.org), to generate standard genomic position format files (BAM) (Langmead and Salzberg, 2012). A summary of the sequencing and alignment statistics is given in **Data S2**. The BAM files were imported into SeqMonk version 1.40.0 (Braham Bioinformatics) to build normalized ChIP-Seq sequence read profiles. To obtain a global view of the binding signals, we subdivided the genome into 1000 bp probes and then calculated the number of reads per probe relative to the total number of reads (Reads per Million; using the Read Count Quantitation option). For each of the different HA-ParB variants, ChIP-seq peaks relative to the total input DNA samples were called with MACS2 v2.1.1.20160309.6 (broad region option), available through the Galaxy analysis platform. The q-value (false discovery rate, FDR) cut-off for called peaks was 0.05. A summary of the peaks identified, rank-ordered according to their location on the chromosome, is provided in **Data S2**. The ChIP-seq profiles in **Figure 4A** show unnormalized read counts based on 10 bp probes. In this case, normalisation was not required as we did not compare reads from different chromosomal segments but only from the *oriC* region.

#### Molecular dynamics simulations

All molecular dynamics (MD) simulations were carried out with the GROMACS program package, version 2019.2 (Abraham et al., 2015). The starting structures for the MD simulations were generated from the X-ray crystal structures of the *Mx*ParB·CDP·P_S_, *Mx*ParB_Q52A_·CDPγS and *Mx*ParB_E93A_·CDP·P_S_ complexes reported in this study. Since our MD simulations focus on the pre-hydrolysis state of ParB, CDP+P_s_ were converted to CTP in the wild-type and E93A structures by superimposing the coordinates of the triphosphate-Mg^2+^ moiety of CTPγS taken from the X-ray structure of the Q52A variant after conversion of CTPγS to CTP.

The Amber ff99SB-ILDN (Hornak et al., 2006; Lindorff-Larsen et al., 2010) force field was used for the protein and the force field parameters of Meagher and coworkers (Meagher et al., 2003) were used for the nucleotide. The systems were solvated with SPC/E (Berendsen et al., 1987) water. All crystal water molecules were maintained in the simulation setup. MD simulations were carried out with periodic boundary conditions using cubic simulation boxes with ca. 9.6 nm edge length, which contained a total of ca. 28,000 atoms including water molecules and Na^+^ ions to neutralize the system. Prior to the simulations, the systems were energy-minimized and equilibrated for 5 ns in the NpT ensemble with harmonic position restraints on all protein and CTP heavy atoms (force constants of 1000 kJ/mol/nm^2^) to relax the water around the protein. After this equilibration, for each of the three different systems (WT, E93A, Q52A), five 200-ns production simulations were carried out in the NpT ensemble at 300 K, initiated using different seeds to generate the initial atomic velocities from a Maxwell-Boltzmann distribution. The total sampling time of the production simulations thus amounts to 3000 ns. However, since ParB is a homodimer, each simulation samples two nucleotide-binding sites.

Temperature was kept constant during the MD simulations by coupling to the velocity rescaling thermostat of Bussi and coworkers (Bussi et al., 2007) with a coupling time constant of 0.1 ps. Constant 1 bar pressure was maintained using a weak coupling barostat (Berendsen et al., 1984) with a time constant of 2 ps and a compressibility of 4.5 × 10 bar^-1^. Short-range non-bonded Coulomb and Lennard-Jones 6,12 interactions were treated with a buffered Verlet pair list (Páll and Hess, 2013) with potentials smoothly shifted to zero at a 1.0 nm cutoff. Analytical dispersion corrections for energy and pressure were applied to compensate for the truncation of the Lennard-Jones interactions. Long-range electrostatic interactions were treated with the particle mesh Ewald (PME) (Darden et al., 1993) method with default settings. The LINCS (Hess, 2008) and SETTLE (Miyamoto and Kollman, 1992) constraint algorithms were used to constrain all protein bonds involving H atoms and all internal degrees of freedom of water molecules, respectively, allowing to integrate the equations of motion with 2 fs time steps.

#### Stochastic simulation of *Mx*ParB sliding

We developed a stochastic simulation, using the Gillespie algorithm (Gillespie, 1976) and written in C++, for the loading, sliding (diffusion) and dissociation of ParB dimers on the DNA. The DNA of the centromeric region is modelled as a 1D lattice with each lattice site corresponding to 10 bp. We use a lattice with 4000 sites for wild-type ParB and 19,000 sites for the Q52A and E93A variants (corresponding to 40 kb and 190 kb respectively). ParB dimers load onto the DNA at any of the 24 *parS* sites. Loading can only occur if the lattice site is free. Dimers diffuse along the DNA with diffusion coefficient *D* by moving randomly to any neighbouring unoccupied lattice site. Dissociation occurs randomly at a rate *d*. The total number of dimers is fixed at 300. This is motivated by the number estimated for *C. crescentus* (Lim et al., 2014). Any unbound dimers are assumed to be in the cytoplasm, which is well-mixed. The loading rate from the cytoplasm is chosen for every choice of the other parameters such that ∼80% of wild-type dimers are bound at steady state, consistent with the bright foci observed using fluorescence microscopy. All but one of the *parS* sites in *M. xanthus* are located close together in a cluster. While the main peak of the ChIP-seq profile is centred on this cluster, there is a small peak coincident with the isolated *parS* site. This is detectable even after smoothening. However, when we took each *parS* site to have the same loading rate, this small peak was not accurately captured. We therefore increased the relative strength (three-fold) of this site compared to the others such that we obtain the same relative height of this small peak compared to the main peak. This has no effect on the subsequent fitting to the profiles of the Q52A and E93A variants, as their profiles are much wider than the *parS-*containing region.

For each parameter set the simulation is first run until the steady-state is reached. Then the distribution of ParB dimers is recorded at regular time intervals, sufficiently separated to be independent samples of the equilibrium distribution. Before comparing this distribution to the experimental ChIP-seq profiles, we first binned the latter at 10 bp resolution to match the simulations. As the signal is still very noisy, we also smoothen it using the *smooth* function of MATLAB (MathWorks, USA) with a moving window of width 151 (1510 kb). As well as inherent noise and sequence-specific effects, this helps to mitigate the effects of transcription and other roadblocks, which are not implemented in the model. The simulated profile was normalised to have the same area under the curve as the smoothened ChIP-seq profile. We then calculated the mean-squared error between the two profiles (using the MATLAB function *immse)*. The best fit of the simulations to the ChIP-seq data is defined by the parameter *D* and *d* (whichever was being varied) that results in the lowest mean-squared error.

Although the lattice is single-occupancy (sliding ParB dimers cannot move through each other), we found that the simulated profile depends only on the ratio of *D* and *d*, i.e. on the length-scale

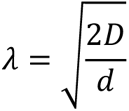

as would be the case for a continuum multi-occupancy model. However, the effect of the single occupancy of the lattice does play a role, since changing the diffusion coefficient can change the steady-state number of dimers bound, depending on the parameters.

The simulations for the wild-type protein were fitted to the ChIP-seq data by varying the diffusion coefficient for a given dissociation rate. Based on our FRAP measurements, we fixed the dissociation rate at *d* = *log*(2)/20 s^-1^.

The ChIP-seq data for the Q52A and E93A variants were fitted by varying the dissociation rate but keeping the diffusion coefficient fixed at the value obtained for wild-type simulations (1.80 x 10^-2^ μm^2^ s^-1^) and using the same loading rate to find the best fit. Parameter fitting was done using the MATLAB function *isqcurvefit*, and 95% confidence intervals were obtained with *nlparci*.

## QUANTIFICATION AND STATISTICAL ANALYSIS

Details on the number of replicates, the sample sizes as well as the value and meaning of n are included in the figure legends. Standard deviations were calculated in Microsoft Excel 2019. Unless indicated otherwise, all experiments were performed at least twice.

## DATA AND CODE AVAILABILITY

The crystallographic data obtained in this study were deposited at the Protein Data Bank (www.rcsb.org) under the accession codes 7BNK (*Mx*ParB·CDP·P_S_), 7BNR (*Mx*ParB_Q52A_·CTPγS) and 7O0N (*Mx*ParB_E93A_·CDP·P_S_). ChIP-seq raw data are available at the Gene Expression Omnibus (GEO) database under the accession number xxx. All other data supporting the findings of this study are included in the main text or the supplemental material. Additional information is available from the authors upon reasonable request.

## SUPPLEMENTAL FIGURES

**Figure S1.**
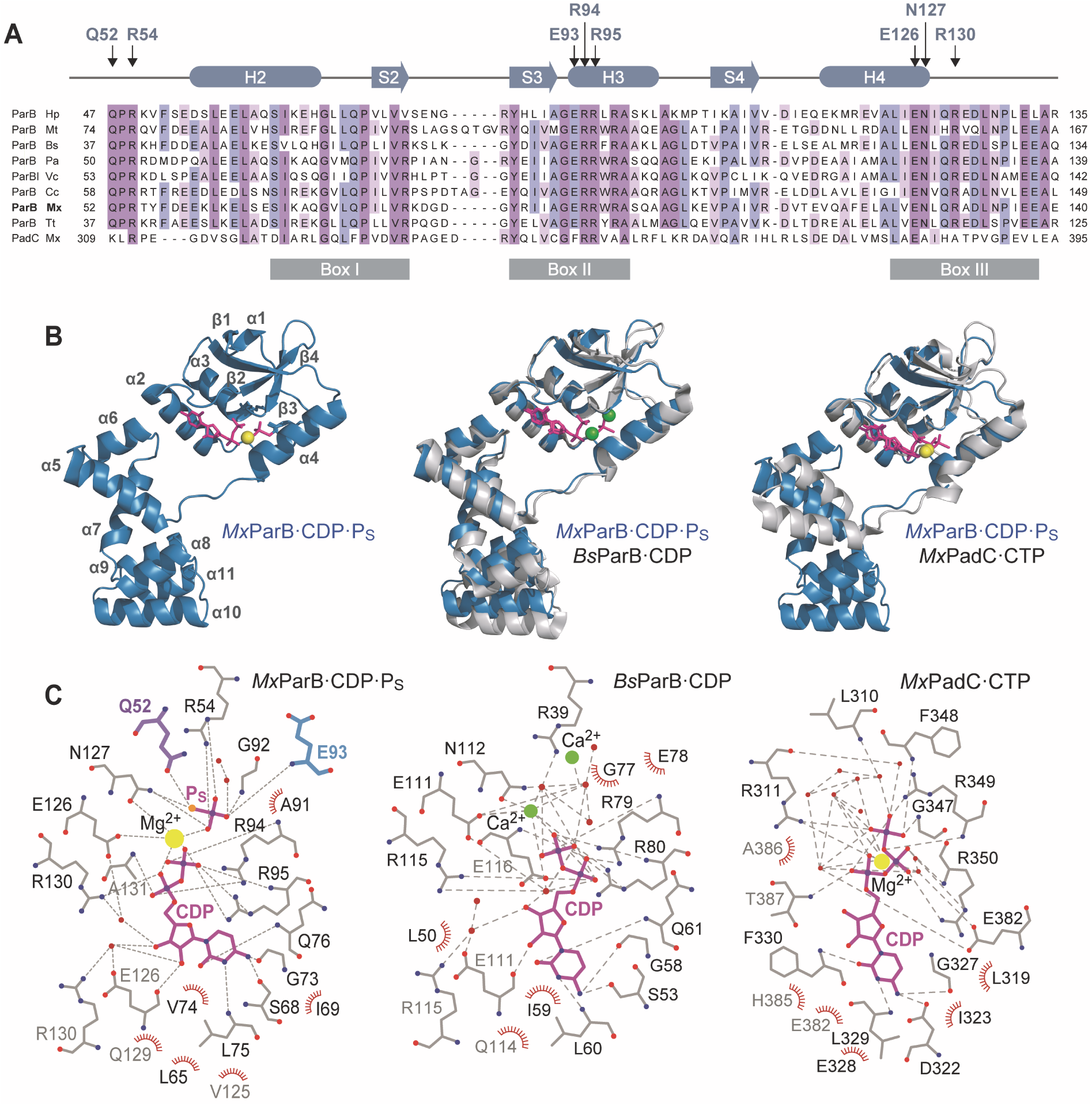
Conservation of the CTP-binding pocket in ParB homologues. Related to Figure 1. **(A)** Amino acid sequence alignment of the CTP-binding region of well-studied ParB homologues. The scheme on top shows the secondary structure elements of *Mx*ParB. Arrows indicate conserved residues in the CTP-binding pocket. The reviously described conserved Box I, Box II and Box III regions (Bartosik et al., 2004; Osorio-Valeriano et al., 2019; Yamaichi et al., 2012) are represented by gray boxes at the bottom. The UniProt accession numbers of the sequences used in the alignment are as follows: O25758 (ParB Hp, *Helicobacter pylori*), P9WIJ9 (ParB Mt, *Mycobacterium tuberculosis*), P26497 (ParB Bs, *Bacillus subtilis*), (ParB Pa, *Pseudomonas aeruginosa*), Q9KNG7 (ParBI Vc, *Vibrio cholerae*), P0CAV8 (ParB Cc, *Caulobacter crescentus*), Q1CVJ4 (ParB Mx, *Myxococcus xanthus*), Q72H91 (ParB Tt, *Thermus thermophilus*), Q1D3H3 (PadC Mx, *Myxococcus xanthus*). **(B)** Arrangement of secondary structural elements in one subunit of the *Mx*ParB·CDP·P_s_ complex in ribbon representation. The CDP and P_S_ molecules are shown in magenta. The Mg^2+^ ion is colored yellow. Structural comparison of ParB·CDP·P_S_ from *M. xanthus* with ParB·CDP from *B. subtilis* (PDB: 6SDK; RMSD of 0.893 Å for 171 paired C_α_ atoms) and PadC·CTP from *M. xanthus* (PDB: 6RYK; RMSD of 0.725 Å for 68 paired C_α_ atoms). **(C)** 2D protein-ligand interaction maps of the different structures shown in (B). The color scheme follows that of Figure 1E.

**Figure S2.**
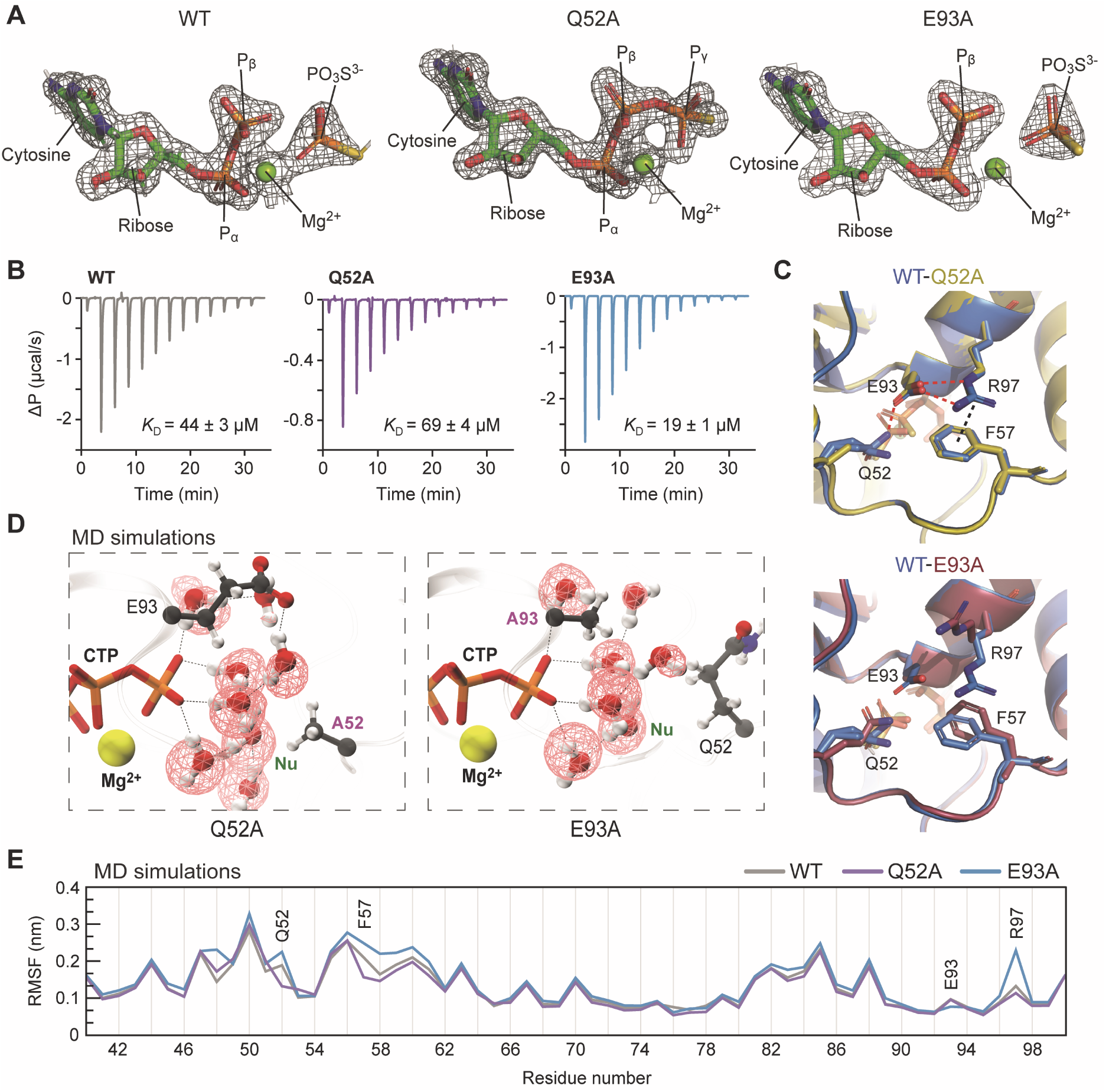
Biochemical and structural characterization of ParB catalytically inactive variants Q52A and E93A. Related to Figure 1. **(A)** Unbiased 2F_obs_-F_calc_ difference electron densities of the cytidine nucleotides (represented as sticks) in the different crystal structures contoured at 2.0 σ and depicted as a gray mesh. Note that the nucleotides were not present during refinement and are only shown for illustration proposes. **(B)** Analysis of the interaction of different ParB variants with CTPγS by isothermal titration calorimetry (ITC). A solution of protein (150 μM) was titrated with 13 consecutive injections (2 μL) of a CTPγS stock solution (1.5 mM). The thermograms show the heat changes observed after each injection. The *K*_D_ values were obtained by fitting the data to a one-set-of-sites model. **(C)** Main structural differences between wild-type ParB and the catalytically inactive Q52A and E93A variants. In the wild-type structure, residue E93 is engaged in hydrogen-bonding interactions with residues Q52 and R97. In the absence of E93, residues Q52, F57 and R97 adopt different conformations. **(D)** Snapshot from MD simulations showing the catalytic site of the Q52A and E93A variants in the CTP-bound, pre-hydrolysis state. Only CTP, the Mg^2+^ ion, the water molecules and residues Q52 (A52) and E93 (A93) are shown for simplicity. The water occupancies are shown as red wire frames. Hydrogen bonds are indicated by red dotted lines. The putative nucleophilic water is labeled in green (Nu). **(E)** Root-mean-square fluctuation (RMSF) plots showing the flexibility of residues 39 to 120 in MD simulations of the indicated *Mx*ParB variants.

**Figure S3.**
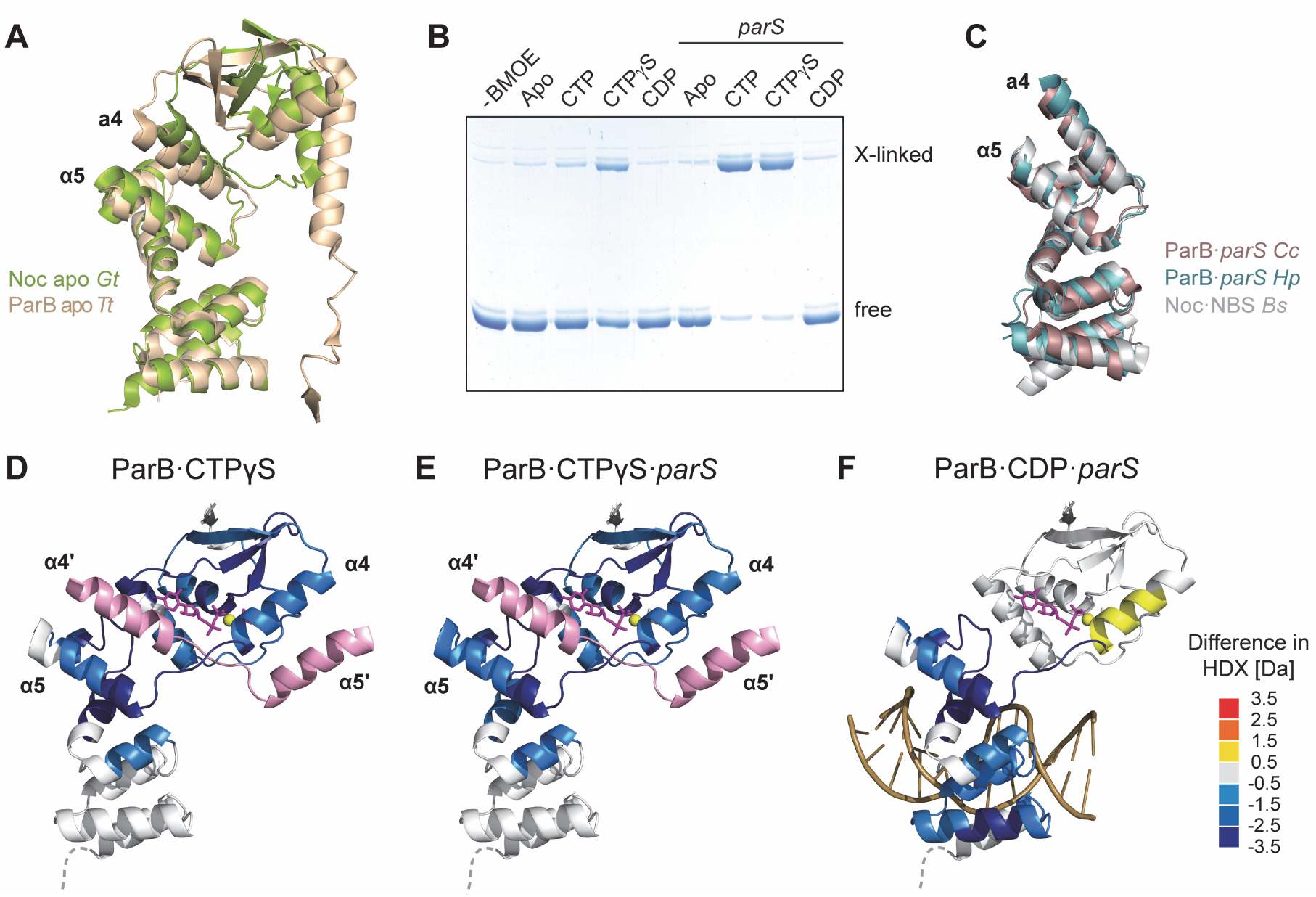
HDX analysis of conformational changes induced in ParB upon ligand binding. Related to Figure 2. **(A)** Structural comparison of apo ParB from *T. thermophilus* (PDB: 1VZ0) and the apo form of the ParB homolog Noc from *Geobacillus thermoleovorans* (PDB: 7NFU). Helices α4 and α5 are labeled. Note that helix α4 bundles with helices α5 and α6 in the DNA-binding domain. **(B)** Coomassie stained SDS-gel showing the cross-linked (X-linked) and non-crosslinked (free) *Mx*ParB-G37C species quantified in Figure 2E. ParB-G37C was incubated with the indicated nucleotide (1 mM) in the absence or presence of *parS* DNA prior to crosslinking with BMOE and separation of the proteins by SDS-PAGE. **(C)** Structural comparison of the DNA-binding domains of ParB homologs crystallized in the presence of their cognate DNA ligand. Shown are the structures of *H. pylori* ParB (PDB: 4UMK) and *C. crescentus* ParB (PDB: 6T1F) in complex with *parS* and *B. subtilis* Noc in complex with the Noc-binding site (NBS) (PDB: 6Y93). The DNA molecules were removed for clarity. **(D and E)** Differences in HDX between apo ParB and ParB·CTPγS in the (D) absence or (E) presence of *parS* DNA, mapped onto the structure of one subunit of *Mx*ParB·CDP·P_S_ complex (t = 30 s; **Figure S4)**. Helices α4 and α5 from the *trans*-subunit of the dimer (α4’ and α5’) are shown in pink. **(E)** Differences in HDX between apo ParB and ParB·CDP·*parS*, mapped onto the structure of one subunit of the *Mx*ParB·CDP·P_S_ complex (t = 30 s; **Figure S4)**. A DNA molecule was modeled into the structure using the crystal structure of the *H. pylori* ParB·*parS* complex (PDB: 4UMK) as a template. The color code is giving on the left.

**Figure S4.**
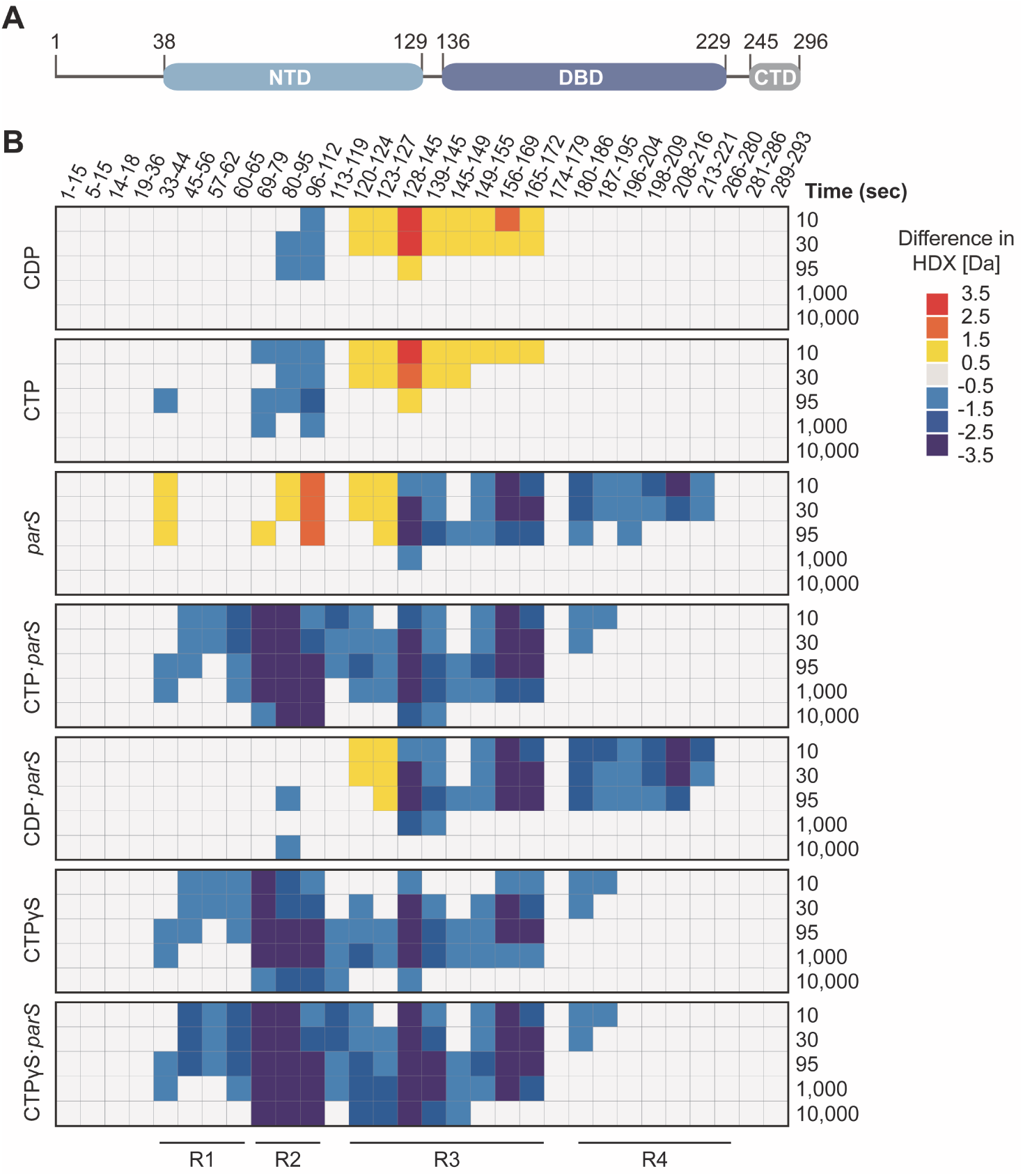
Heat plots of the differences in deuterium uptake between full-length apo ParB and ParB in different ligand-bound states. Related to Figure 2. The heat plot show the differences in deuterium uptake between *Mx*ParB incubated with the indicated nucleotide in the presence or absence of *parS* DNA and apo *Mx*ParB at different incubation times for a series of representative peptides (see **Data S1** for the full set of peptides). The color code is given at the bottom right. The scheme at the top shows the domain organization of ParB. Residues 1-37 comprise the ParA-interacting region, residues 38-129 correspond to the N-terminal CTP-binding domain (NTD), residues 136-229 correspond to the DNA binding domain (DBD) and residues 245-296 to the C-terminal dimerization domain (CTD).

**Figure S5.**
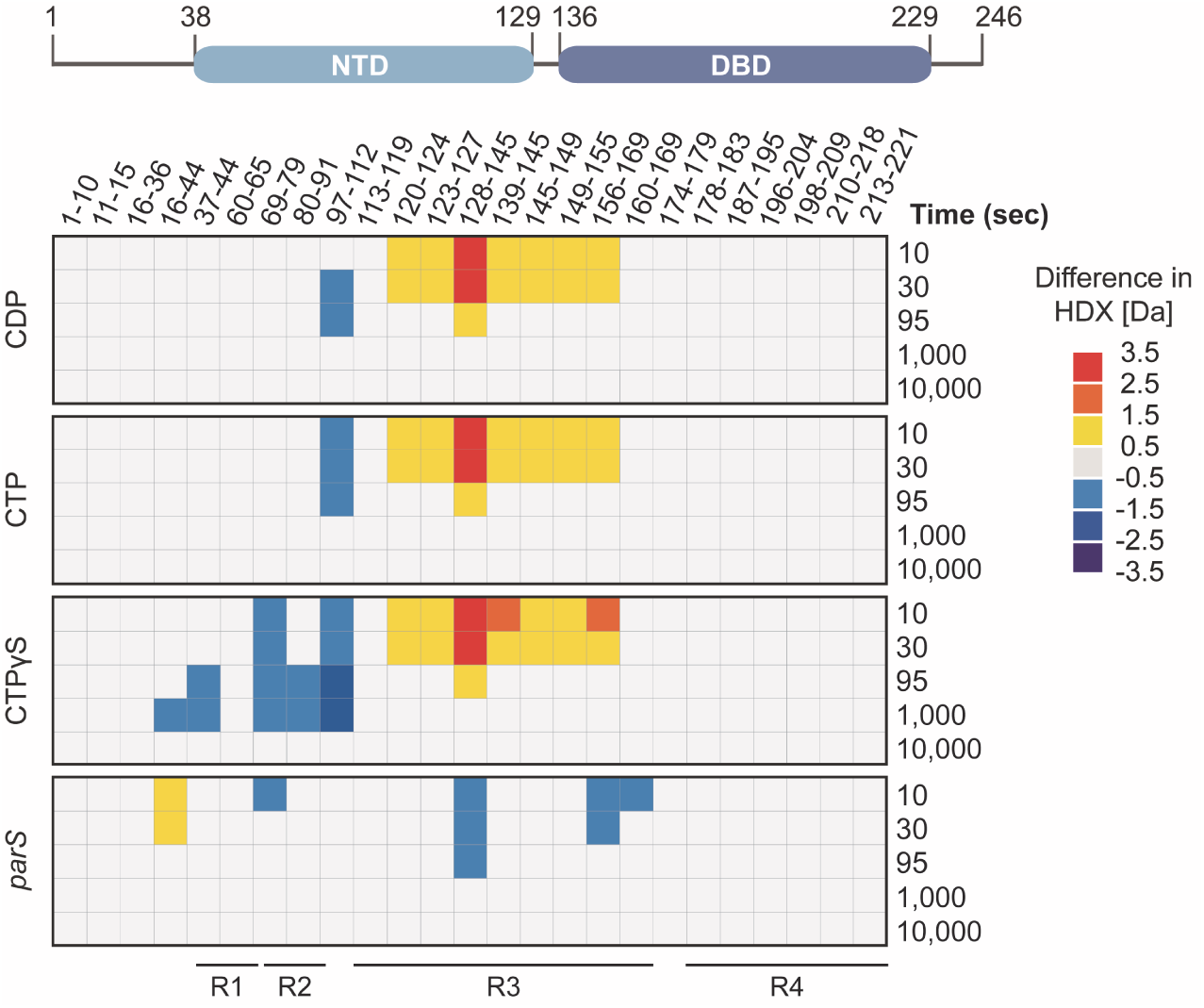
Heat plots of the differences in deuterium uptake between apo ParB_ΔC_ and ParB_ΔC_ in different ligand-bound states. Related to Figure 2. The heat plot shows the differences in deuterium uptake between *Mx*ParB_ΔC_ incubated with the indicated nucleotide in the presence absence of *parS* DNA and apo *Mx*ParB_ΔC_ at different incubation times for a series of representative peptides (see **Data S1** for the full set of peptides). The color code is given at the bottom right. The scheme at the top shows the domain organization of ParB_ΔC_.

**Figure S6.**
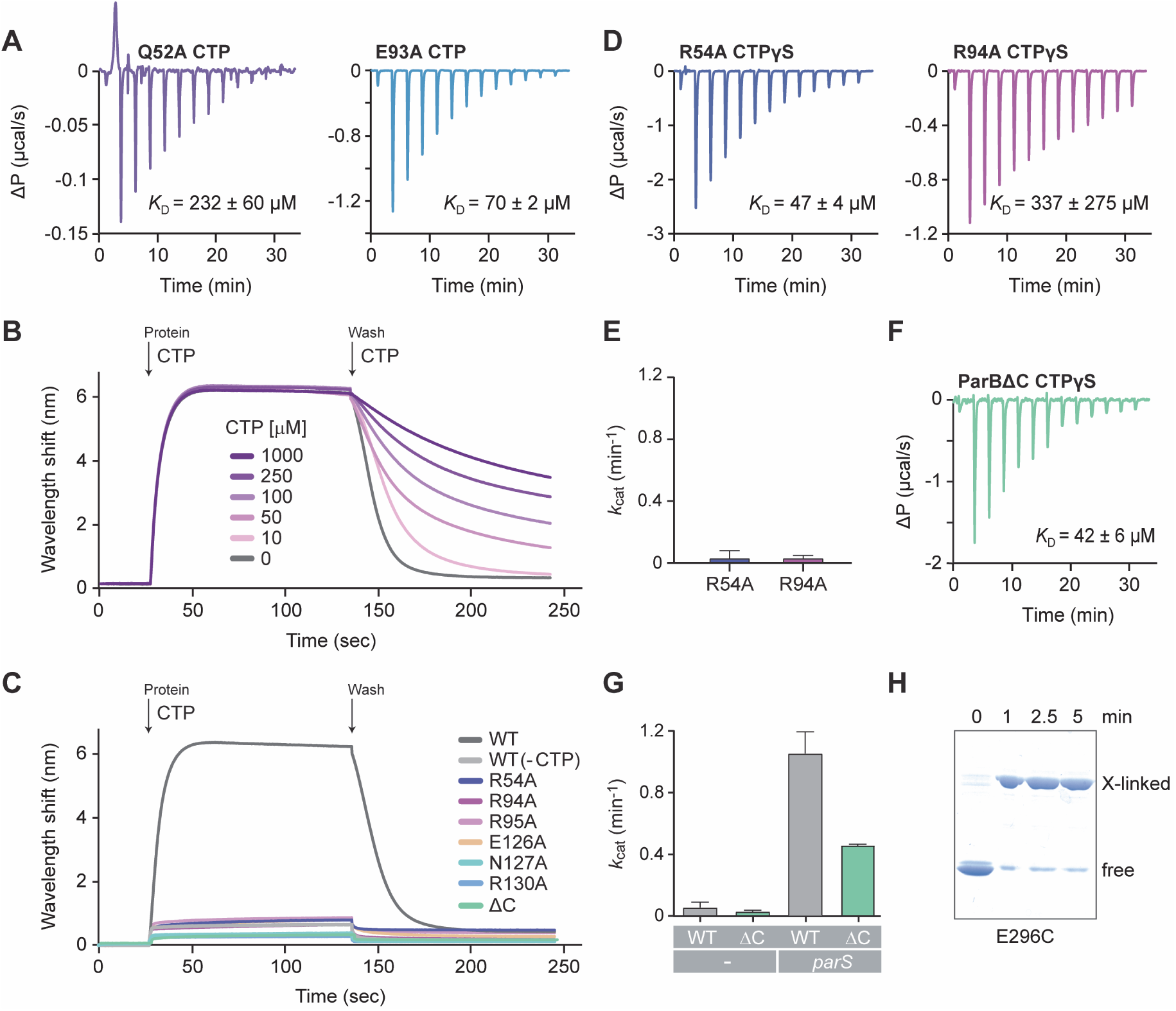
Biochemical analysis of ParB variants. Related to Figure 3. **(A)** Analysis of the interaction of the Q52A and E93A variants with CTP by isothermal titration calorimetry (ITC). A solution of protein (150 μM) was titrated with 13 consecutive injections (2 μL) of a CTP solution (1.5 mM). The thermograms show the heat changes observed after each injection. The corresponding *K*_D_ values were obtained by fitting of the data to a one-set-of-sites model. **(B)** BLI analysis of the unleading rates of *Mx*ParB rings in the presence of varying CTP concentrations. Sensors derivatized with closed dsDNA fragments containing a single *parS* site were loaded with wild-type ParB (10 µM) in the presence of 1 mM CTP. Subsequently, the sensors were transferred to 400 µl of protein-free buffer containing the indicated concentrations of CTP to follow the dissociation of the *Mx*ParB rings. The kinetics of the loading and unloading processes were followed by monitoring the wavelength shifts induced by changes in the optical thickness of the sensor surface during the association and dissociation phases. **(C)** BLI analysis of the accumulation of different ParB variants on *parS*-containing DNA. Sensors derivatized with closed dsDNA substrates were probed with the indicated ParB variants (10 µM) in the presence of CTP (1 mM). To follow the dissociation of ParB rings from DNA, the sensors were transferred into 400 µL of protein- and nucleotide-free buffer. **(D)** Analysis of the interaction of the R54A and R94A variants with CTPγS by isothermal titration calorimetry (ITC), performed as in (A). **(E)** CTPase activities of the *Mx*ParB R54A and R94A variants. The indicated proteins (5 µM) were incubated with 1 mM CTP in the presence of *parS* DNA (250 nM). The reaction rates were determined with an NADH-coupled enzyme assay. Data represent the mean of three replicates (± SD). **(F)** Analysis of the interaction of ParB_ΔC_ with CTPγS by isothermal titration calorimetry (ITC), performed as in (A). **(G)** CTPase activities of wild-type *Mx*ParB and *Mx*ParB_ΔC_ in the presence or in the absence of *parS* DNA. **(H)** Coomassie stained SDS-gel showing the crosslinked (X-linked) and non-crosslinked (free) species of *Mx*ParB-E296C after BMOE treatment. The protein (10 µM) was incubated with BMOE (1 mM). Samples were taken at the indicated time points and analyzed by SDS-PAGE.

**Figure S7.**
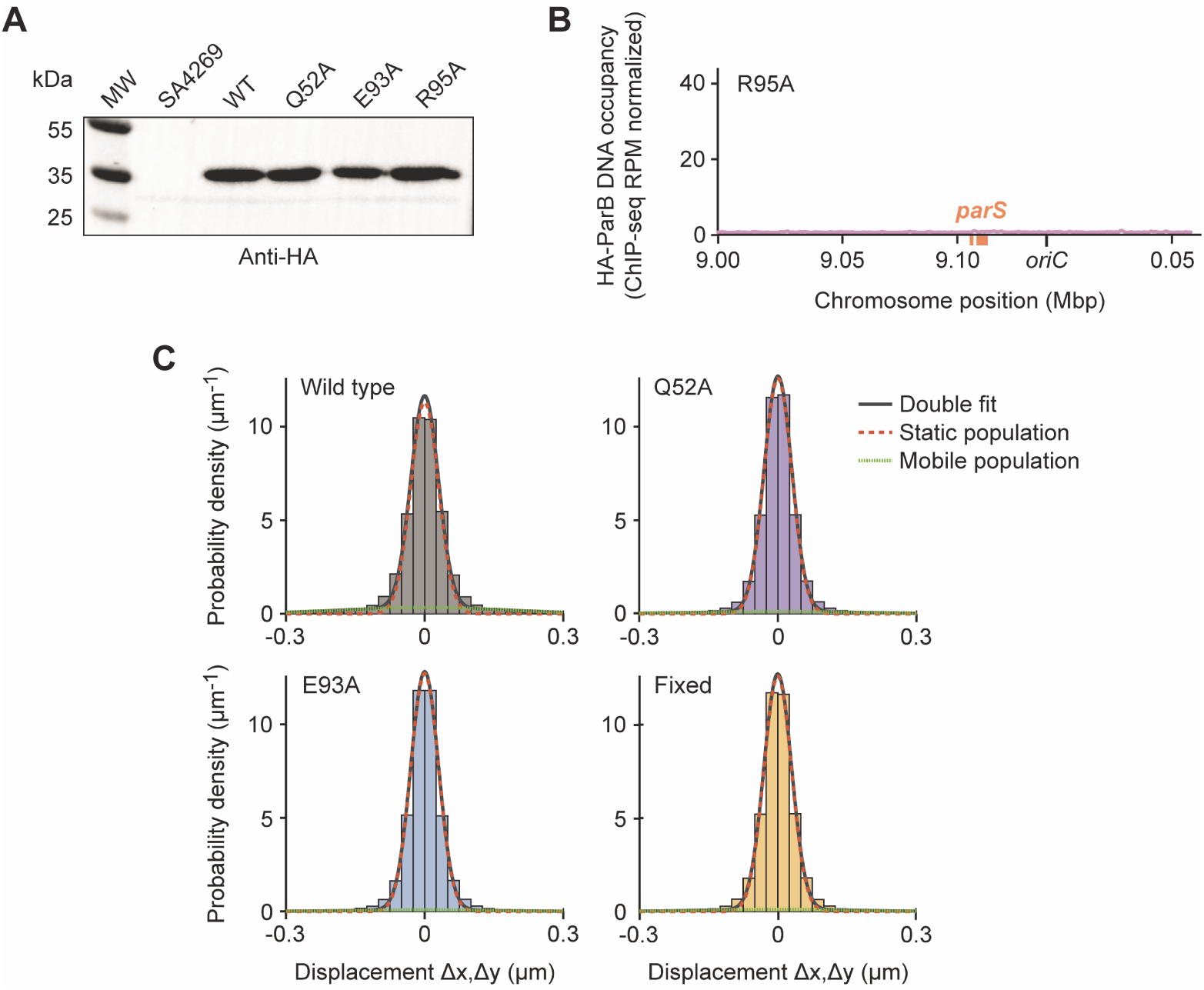
*In vivo* analysis of ParB mutant variants. Related to Figures 4 and 5. **(A)** Western blot analysis of strains producing the different HA-ParB variants used for the ChIP-seq analysis. Cultures were treated as described in Figure 4A prior to Western blot analysis with anti-HA antibodies. A molecular weight marker (M) was applied as a reference. The background strain SA4269 (Δ*parB* P_cuoA_-*parB*) was analyzed as a negative control. ChIP-seq analysis of a CTP-binding-deficient HA-ParB variant (R95A). *M. xanthus* strain MO84 (R95A) was grown as described in Figure 4A prior to ChIP-seq analysis. Normalized counts in reads per million (rpm) are plotted against the chromosomal position for an ∼200 kb region centered at *oriC*. **(C)** Gaussian mixture model (GMM) analysis of the mobility of the different mNeonGreen-*Mx*ParB variants shown in Figure 5B. The probability distributions of the frame-to-frame displacements in both x and y direction from single-particle tracking experiments were fitted to a two-component Gaussian function (sum in black), assuming a fast-diffusing mobile (green) and a slow-diffusing (orange) population.

## SUPPLEMENTAL TABLES

**Table S1.** Crystallographic data collection and refinement statistics.

|  | *MxParB* | *MxParB<sub>Q52A</sub>* | *MxParB<sub>E93A</sub>* |
| --- | --- | --- | --- |
| **Data collection** |  |  |  |
| Space group | *P*2<sub>1</sub>2<sub>1</sub>2<sub>1</sub> | *P*2<sub>1</sub>2<sub>1</sub>2<sub>1</sub> | *P*2<sub>1</sub>2<sub>1</sub>2<sub>1</sub> |
| Cell dimensions |  |  |  |
| *a*, *b*, *c* (Å) | 44.243 80.706 143.38 | 45.75 80.87 135.71 | 44.12 80.37 136.83 |
| *a*, *b*, *c* (°) | 90 90 90 | 90 90 90 | 90 90 90 |
| Resolution (Å) | 42.28 - 1.9 (1.968 - 1.9) | 40.44 - 1.501 (1.555 - 1.501) | 41.99 - 1.89 (1.958 - 1.89) |
| *R*<sub>merge</sub> | 0.1961 (2.221) | 0.06803 (2.329) | 0.09372 (1.938) |
| *I* / σ*I* | 9.21 (0.83) | 19.67 (0.74) | 14.18 (1.09) |
| Completeness (%) | 99.87 (99.61) | 99.82 (98.85) | 99.63 (96.81) |
| Redundancy | 7.3 (7.4) | 12.8 (9.7) | 12.3 (11.4) |
| CC<sub>1/2</sub> | 0.996 (0.457) | 1 (0.457) | 0.999 (0.578) |
| **Refinement** |  |  |  |
| Resolution (Å) | 42.28 – 1.90 | 40.44 – 1.70 | 44.12 – 1.89 |
| No. reflections | 41418 (4081) | 56287 (5557) | 39707 (3794) |
| *R*<sub>work</sub> / *R*<sub>free</sub> | 20.67/24.38 | 18.23/21.07 | 19.55/21.65 |
| No. atoms |  |  |  |
| Protein | 30432 | 3076 | 3079 |
| Ligand/ion | 87 | 66 | 81 |
| Water | 159 | 306 | 60 |
| *B*-factors |  |  |  |
| Protein | 46.46 | 39.69 | 34.54 |
| Ligand/ion | 39.55 | 25.81 | 46.48 |
| Water | 46.19 | 39.27 | 51.62 |
| Ramachandran (%) |  |  |  |
| favored | 98.43 | 98.71 | 98.70 |
| allowed | 1.57 | 1.29 | 1.30 |
| outliers | 0.00 | 0.00 | 0.00 |
| R.m.s. deviations |  |  |  |
| Bond lengths (Å) | 0.005 | 0.005 | 0.010 |
| Bond angles (°) | 0.88 | 1.02 | 1.32 |

\*Values in parentheses are for highest-resolution shell.

**Table S2.** Characteristics of mutant ParB variants.

| Variant | CTPase activity<br>$k_{cat}$ (min<sup>-1</sup>) | CTP binding<br>$K_D$ (μM) | Localization<br>pattern | BLI<br>Signal, dissociation | ChIP-seq<br>Region covered (kb) |
| --- | --- | --- | --- | --- | --- |
| Wild type | 1.0 | 59 | foci | high | 33 |
| R54A | ND | 47 | - | low | - |
| Q52A | ND | 69 | foci | high, slow dissociation | 175 |
| E93A | 0.1 | 19 | foci | high, slow dissociation | 125 |
| R94A | ND | 337 | - | low | - |
| R95A | ND<sup>1</sup> | ND<sup>1</sup> | diffuse<sup>1</sup> | low | ND |
| E126A | 0.1<sup>1</sup> | 31<sup>1</sup> | speckles<sup>1</sup> | low | - |
| N127A | ND<sup>1</sup> | 137<sup>1</sup> | diffuse<sup>1</sup> | low | - |
| R130A | ND<sup>1</sup> | ND<sup>1</sup> | diffuse<sup>1</sup> | low | - |

ND: not detectable
“-”: not determined
<sup>1</sup> Osorio-Valeriano *et al.* (2019)

**Table S3.**
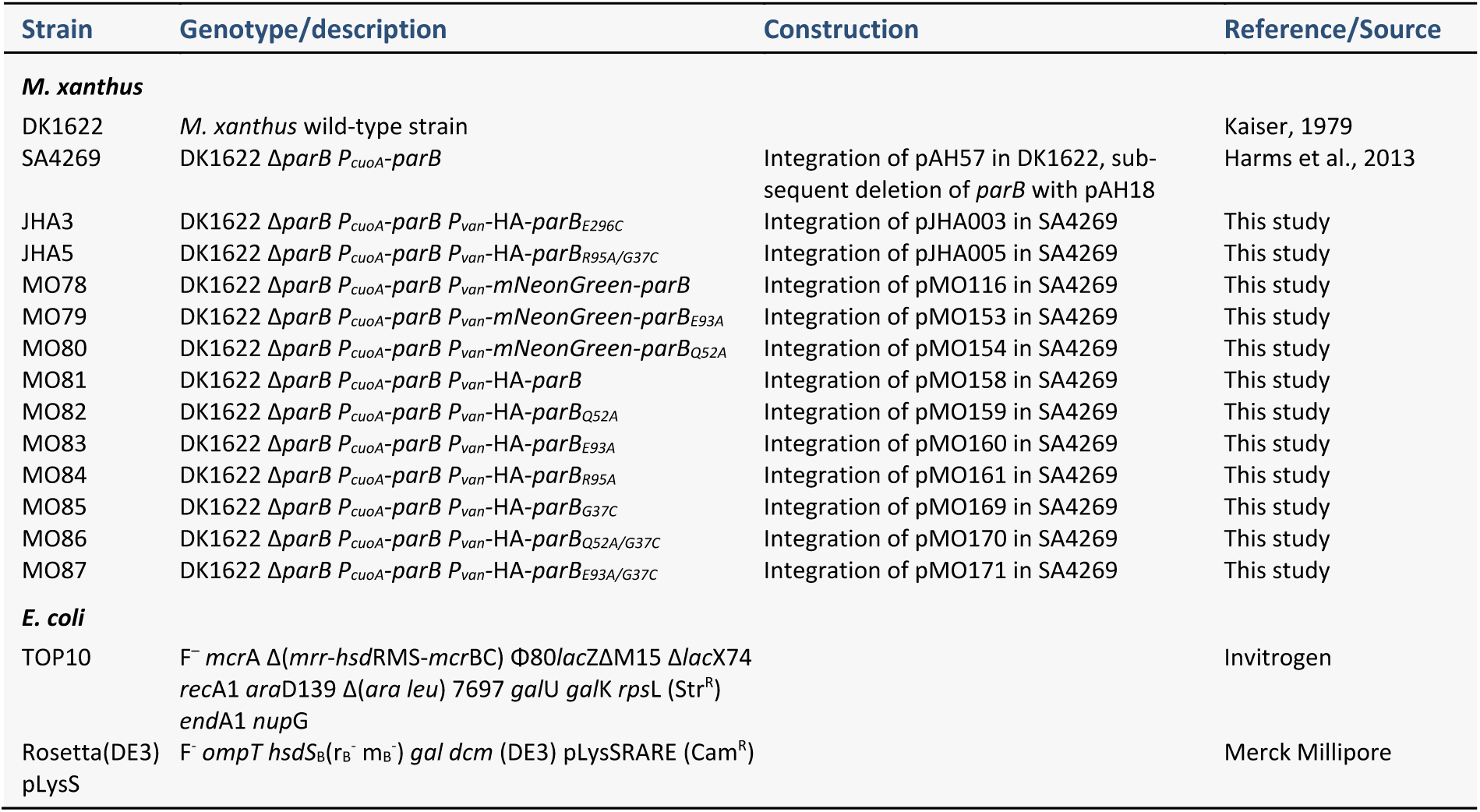
Strains used in this study.

**Table S4.** Plasmids generated in this work.

| Plasmid | Description | Construction |
| --- | --- | --- |
| pJHA3 | pMR3690 bearing HA-*parB*<sub>E296C</sub> | a) PCR amplification of HA-*parB*<sub>E296C</sub> from pMO166 with primers MO258 and M259<br>b) The HA-*parB*<sub>E296C</sub> fragment was used as template for a second PCR reaction with primers MO261 and MO266<br>c) Insertion of the fragment into pMR3690 cut with NdeI and EcoRI by Gibson assembly |
| pJHA5 | pMR3690 bearing HA-*parB*<sub>R95A/G37C</sub> | a) Site directed mutagenesis of pMO161 with primers MO235 and MO236 |
| pLS3 | pTB146 bearing *parB*<sub>R95A</sub> | Site directed mutagenesis of pMO104 with primers MO175 and MO176 |
| pLS4 | pTB146 bearing *parB*<sub>E126A</sub> | a) PCR amplification of *parB*<sub>E126A</sub> from pMO108 with primers MO169 and MO170<br>b) Insertion of the *parB*<sub>E126A</sub> fragment into pTB146 cut with BamHI and SapI by Gibson assembly |
| pMO104 | pTB146 bearing *parB* | a) PCR amplification of *parB* with primers MO169 and MO170<br>b) Insertion of the *parB* fragment into pTB146 cut with BamHI and SapI by Gibson assembly |
| pMO106 | pTB146 bearing *parB*<sub>ΔC</sub> | a) PCR amplification of *parB*<sub>ΔC</sub> with primers MO169 and MO174<br>b) Insertion of the *parB*<sub>ΔC</sub> fragment into pTB146 cut with BamHI and SapI by Gibson assembly |
| pMO116 | pMR3690 bearing *mNeonGreen-parB* | a) PCR amplification of *mNeonGreen*-L1 with primers MO193 and MO194<br>b) The *mNeonGreen*-L1 fragment was used as template for a second PCR reaction with primers MO196 and MO197<br>c) PCR amplification of *parB* with primers pMO198 and pMO199<br>d) Insertion of the *mNeonGreen*-L1 and *parB* fragments into pMR3690 cut with NdeI and EcoRI by Gibson assembly |
| pMO137 | pTB146 bearing *parB*<sub>ΔNC</sub> | a) PCR amplification of *parB*<sub>ΔNC</sub> with primers MO174 and MO243<br>b) Insertion of the *parB*<sub>ΔNC</sub> fragment into pTB146 cut with BamHI and SapI by Gibson assembly |
| pMO138 | pTB146 bearing *parB*<sub>G37C</sub> | Site-directed mutagenesis of pMO104 with primers MO235 and MO236 |
| pMO139 | pTB146 bearing *parB*<sub>Q52A</sub> | Site-directed mutagenesis of pMO104 with primers MO237 and MO238 |
| pMO141 | pTB146 bearing *parB*<sub>E93A</sub> | Site-directed mutagenesis of pMO104 with primers MO241 and MO242 |
| pMO146 | pTB146 bearing *parB*<sub>R94A</sub> | Site-directed mutagenesis of pMO104 with primers MO244 and MO245 |
| pMO153 | pMR3690 bearing *mNeonGreen-parB*<sub>E93A</sub> | Site-directed mutagenesis of pMO116 with primers MO241 and MO242 |
| pMO154 | pMR3690 bearing *mNeonGreen-parB*<sub>Q52A</sub> | Site-directed mutagenesis of pMO116 with primers MO237 and MO238 |
| pMO155 | pTB146 bearing *parB*<sub>R54A</sub> | Site-directed mutagenesis of pMO104 with primers MO253 and MO254 |
| pMO158 | pMR3690 bearing HA-*parB* | a) PCR amplification of HA-*parB* from pMO104 with primers MO259 and MO260<br>b) The HA-*parB* fragment was used as template for a second PCR reaction with primers MO261 and MO199<br>c) Insertion of the HA-*parB* fragment into pMR3690 cut with NdeI and EcoRI by Gibson assembly |
| pMO159 | pMR3690 bearing HA-*parB*<sub>Q52A</sub> | Site-directed mutagenesis of pMO158 with primers MO237 and MO238 |
| pMO160 | pMR3690 bearing HA-*parB*<sub>E93A</sub> | Site-directed mutagenesis of pMO158 with primers MO241 and MO242 |
| pMO161 | pMR3690 bearing HA-*parB*<sub>R95A</sub> | Site-directed mutagenesis of pMO158 with primers MO175 and MO176 |
| pMO166 | pTB146 bearing *parB*<sub>E296C</sub> | Site-directed mutagenesis of pMO104 with primers MO257 and MO258 |
| pMO167 | pTB146 bearing *parB*<sub>Q52A/G37C</sub> | Site-directed mutagenesis of pMO139 with primers MO235 and MO236 |
| pMO168 | pTB146 bearing *parB*<sub>E93A/G37C</sub> | Site-directed mutagenesis of pMO141 with primers MO235 and MO236 |
| pMO169 | pMR3690 bearing HA-*parB*<sub>G37C</sub> | Site-directed mutagenesis of pMO158 with primers MO235 and MO236 |
| pMO170 | pMR3690 bearing HA-*parB*<sub>Q52A/G37C</sub> | Site-directed mutagenesis of pMO159 with primers MO235 and MO236 |
| pMO171 | pMR3690 bearing HA-*parB*<sub>E93A/G37C</sub> | Site-directed mutagenesis of pMO160 with primers MO235 and MO236 |
| pMO173 | pTB146 bearing *parB*<sub>Q52AΔNC</sub> | a) PCR amplification of *parB*<sub>Q52AΔNC</sub> from pMO139 with primers MO174 and MO243<br>b) Insertion of the *parB*<sub>Q52AΔNC</sub> fragment into pTB146 cut with BamHI and SapI by Gibson assembly |
| pMO174 | pTB146 bearing *parB*<sub>E93AΔNC</sub> | a) PCR amplification of *parB*<sub>E93AΔNC</sub> from pMO141 with primers MO174 and MO243<br>b) Insertion of the *parB*<sub>E93AΔNC</sub> fragment into pTB146 cut with BamHI and SapI by Gibson assembly |
| pMO175 | pTB146 bearing *parB*<sub>E126A</sub> | Site-directed mutagenesis of pMO104 with primers MO179 and MO180 |
| pMO176 | pTB146 bearing *parB*<sub>N127A</sub> | Site-directed mutagenesis of pMO104 with primers MO183 and MO184 |
| pMO177 | pTB146 bearing *parB*<sub>R1307A</sub> | Site-directed mutagenesis of pMO104 with primers MO185 and MO186 |

**Table S5.**
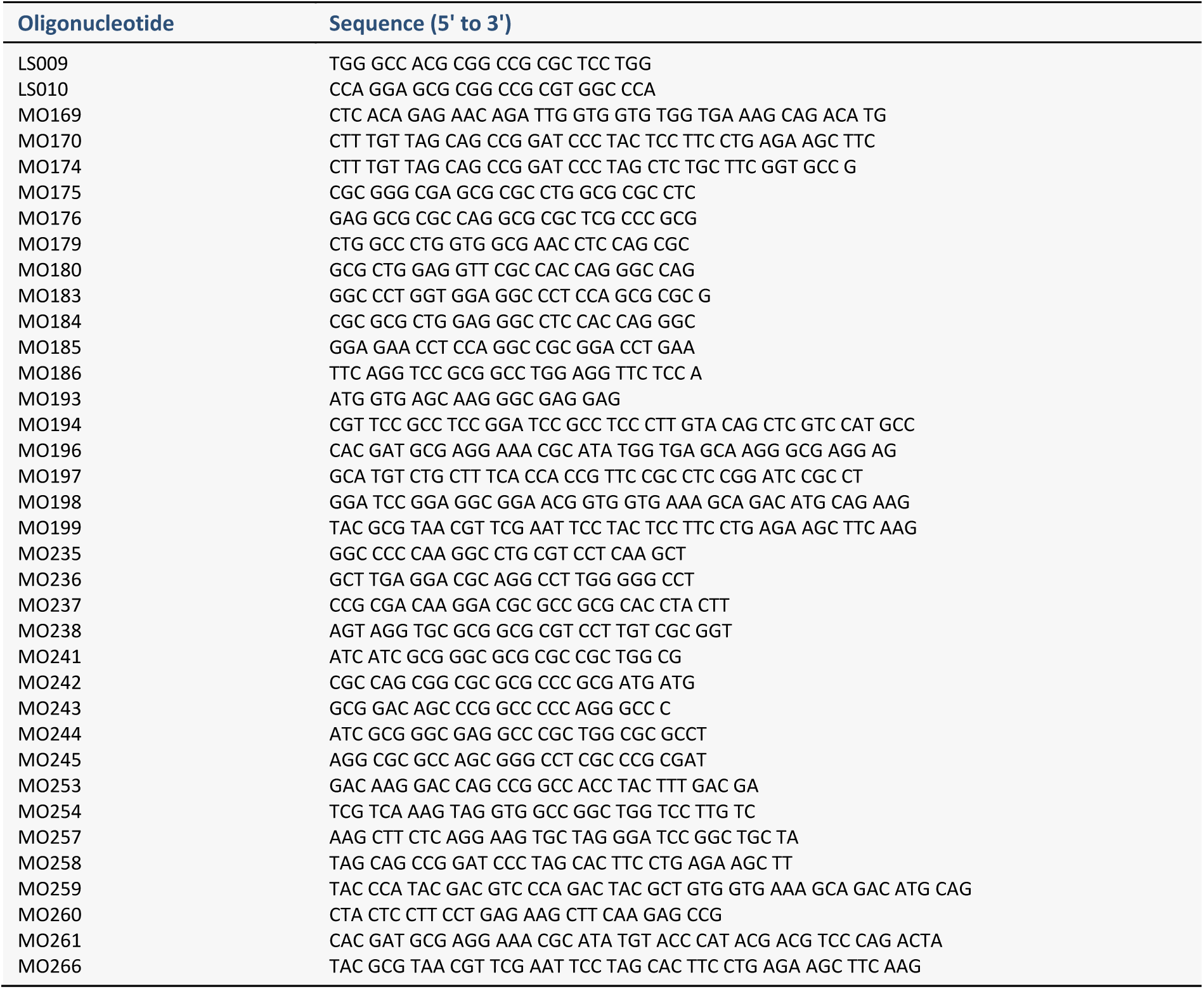
PCR primers used in this work.

